# Bioluminescence-Based Determination of Cytosolic Accumulation of Antibiotics in *Escherichia coli*

**DOI:** 10.1101/2023.12.06.570448

**Authors:** Rachita Dash, Kadie A. Holsinger, Mahendra D. Chordia, Mohammad Sharifian Gh., Marcos M. Pires

## Abstract

Antibiotic resistance is an alarming public health concern that affects millions of individuals across the globe each year. A major challenge in the development of effective antibiotics lies in their limited ability to permeate into cells, noting that numerous susceptible antibiotic targets reside within the bacterial cytosol. Consequently, improving cellular permeability is often a key consideration during antibiotic development, underscoring the need for reliable methods to assess the permeability of molecules across cellular membranes. Currently, methods used to measure permeability often fail to discriminate between arrival within the cytoplasm and the overall association of molecules with the cell. Additionally, these techniques typically possess throughput limitations. In this work, we describe a luciferase-based assay designed for assessing the permeability of molecules into the cytosolic compartment of Gram-negative bacteria. Our findings demonstrate a robust system that can elucidate the kinetics of intracellular antibiotics accumulation in live bacterial cells in real time.

## Introduction

Antibiotic resistance is a growing global problem that is directly leading to increased risks associated with bacterial infections. Recent data reveal that antibiotic resistance was responsible for nearly 5 million fatalities in 2019.^1^ A primary driver of the resistance phenotype is the overuse and misuse of antibiotics in human medicine and agriculture, as well as the lack of development of new antibiotics.^2^ Infections that are highly resistant can lead to prolonged illness, increased healthcare costs, and higher mortality rates. Urgent measures, including responsible antibiotic stewardship, innovative drug development, and public awareness, are essential to combat this pressing threat to modern medicine. Consequently, it is important to prioritize strategies aimed at the circumvention of antibiotic resistance.

Among the primary hurdles in the development of effective antibiotics is their general lack of cellular permeability.^3^ This challenge is particularly pronounced when targeting Gram-negative bacteria and mycobacteria due to their additional membranes that pose barriers to molecular entry. Compounding this issue is the fact that some of the most crucial drug targets are situated within the bacterial cytosol, emphasizing the need for permeable antibiotics in combatting bacterial infections.^4^ In the pursuit of novel antibiotics, it becomes critically important to develop methodologies that can reliably report on molecule accumulation in bacteria with high efficiency.^5–12^ Currently, several methods are available to evaluate the permeability of molecules into Gram-negative bacteria. The most widely used method of LC-MS/MS does not require a chemical tag to be added to the test molecule^13,14^. However, its widespread adoption has been significantly hampered by inherent throughput limitations, limiting its broad application in the field. Another widely used approach involves optical analysis of cells that are treated with compounds chemically modified with a fluorophore to track their entry into the cell.^15,16^ Critically, these methods often fail to report on whether the molecules arrive within the cytoplasmic space and, instead, provide information on the total association of the molecules with the target cells.^17^

Our group has recently described methods to interrogate the accumulation of molecules onto the surface of Gram-positive bacteria and past the outer membrane of diderm bacteria using a combination of click chemistry^18,19^ and HaloTag^20^. Herein, we sought to establish a luciferase-based assay^21^ to determine the accumulation of molecules to the cytosol of Gram-negative bacteria in real time. In this assay, luciferase-expressing bacteria in the presence of 6-hydroxy-2-cyanobenzothiazole (CBT-OH) are incubated with test molecules tagged *via* a disulfide bond to D-Cysteine (D-Cys)^22^ (**Fig. 1a**). Upon the arrival of the conjugate in the reducing environment of the cytosol^23^, reduction of the disulfide bond to generate D-Cys in the cytosol enables a fast and biorthogonal^24^ recombination of CBT and D-Cys to generate intracellular D-luciferin that is rapidly processed by luciferase to generate light (Fig. 1a; **Fig. S1**).^25^ We showed that the system was robust, displayed a high signal-to-noise ratio, and revealed the kinetics of intracellular accumulation of antibiotics.

**Figure 1.**
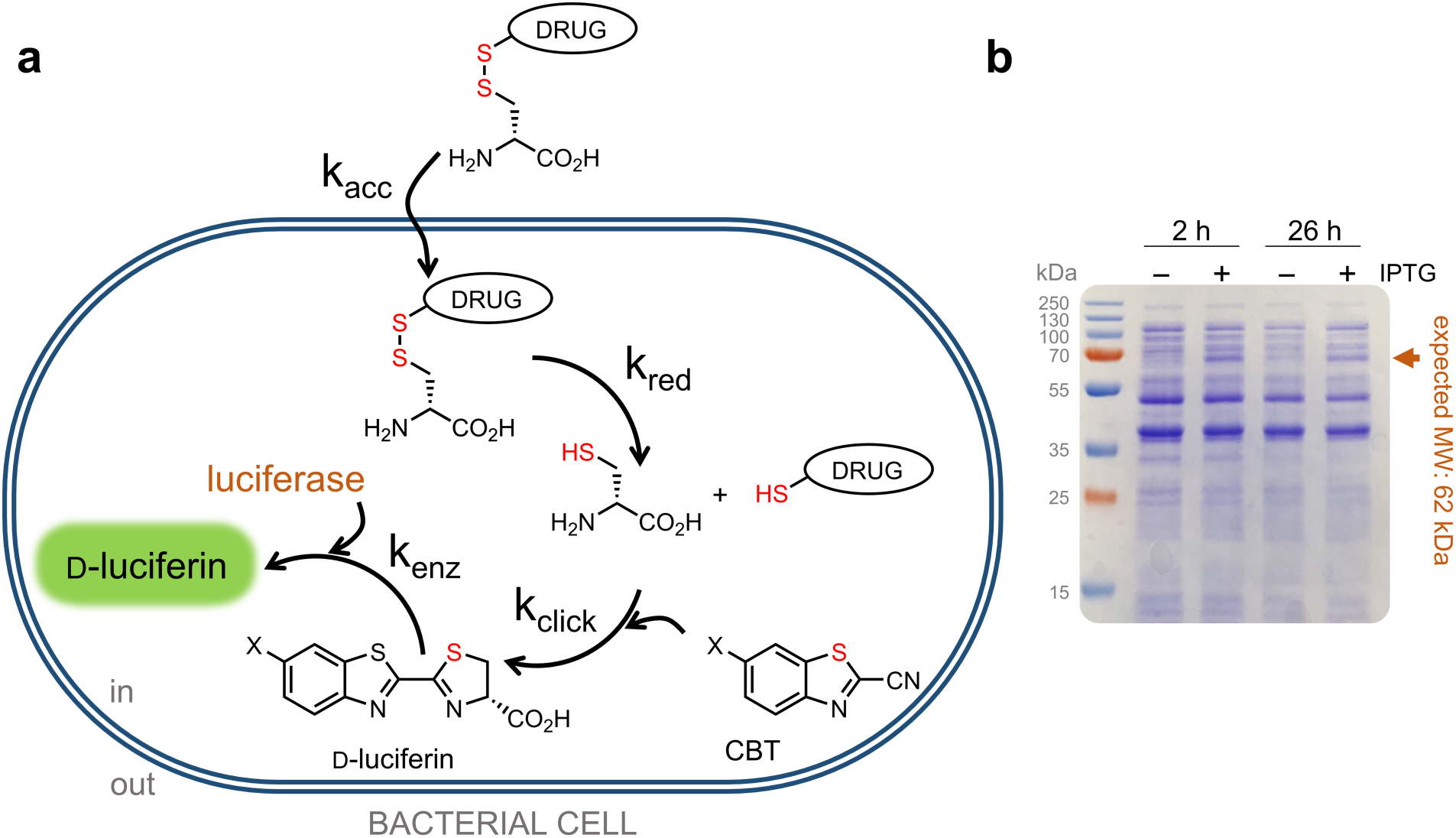
**(a)** Schematic showing the workflow of DCCAA. **(b)** SDS-PAGE analysis of luciferase protein expression in *E. coli* at 2 h or 26 h post IPTG induction. A molecular weight ladder with sizes in kilodaltons (kDa) is shown. The expected molecular weight of luciferase is 62 kDa.

## Results and Discussion

The reaction between aminothiols (including cysteines) and CBT exhibits numerous favorable characteristics, including reaction speed, ease of use, and exceptional specificity. This reaction has been leveraged in a variety of applications including protein labeling^26–28^ and molecular imaging^29^, and to investigate the permeability of peptides into mammalian cells.^21^ Alternatively, mice expressing luciferase have been used to detect endogenous D-Cys in the brain of animals.^30,31^ We identified many advantages to the D-Cys cytosolic accumulation assay (DCCAA). DCCAA generates real time measurement upon singular cellular treatment (no washing steps or chase treatment required), thereby reducing assay manipulation steps. Additionally, the assay workflow is compatible with multi-well plates thus enabling high-throughput analysis. Finally, bioluminescence signals are more compatible with bacterial species that have intrinsic fluorescence, which can introduce high background noise in fluorescence-based assays.

The first step was to evaluate the expression of luciferase in *E. coli*. Bacterial cells transformed with the luciferase-expressing plasmid were grown to mid-log phase and induced with isopropyl-β-D-1-thiogalactopyranoside (IPTG). Two conditions were tested: a 2-h short induction and a 26-h long induction.^32,33^ Both induced and uninduced samples were collected and protein expression was evaluated *via* sodium dodecyl sulfate polyacrylamide gel electrophoresis (SDS-PAGE). The presence of a band consistent with the molecular weight of luciferase (62 kDa) was identified (**Fig. 1b**). Our results showed similar expression levels for both time points, therefore the shorter incubation time (2 h) was selected for subsequent assays.

An assessment of the difference in luminescence signal between induced and uninduced samples in DCCAA was then performed. Bioluminescence imaging of a multi-well plate containing luciferase-expressing *E. coli* cells treated with D-luciferin also revealed a notable increase in luminescence upon induction (**Fig. 2a**). *E. coli* cells harboring the luciferase plasmid were co-incubated with CBT-OH and D-Cystine for one hour, during which luminescence was recorded. Here, D-Cystine served as a surrogate for a test molecule (**Fig. 2b**). Kinetic analysis of the cellular treatment revealed a pronounced increase in the luminescence signal over time. The signal response was IPTG-dependent, which is consistent with the proposed luciferase mediated processing D-luciferin upon generation of D-Cys in the cytosol.

**Figure 2.**
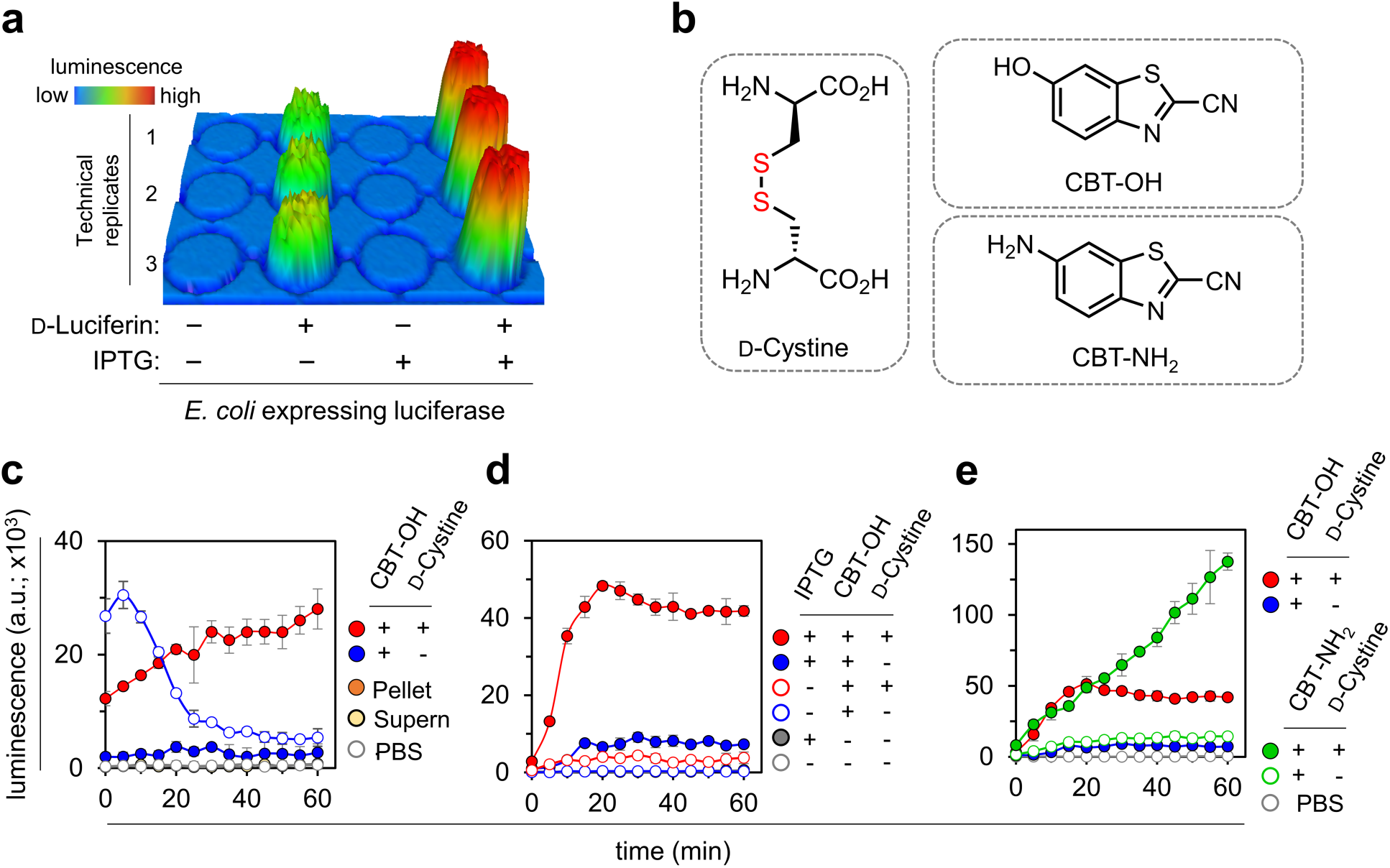
**(a)** Bioluminescence image of a 96-well plate containing *E. coli* treated with100 μM D-luciferin in the presence and absence of IPTG induction. (**c**) Bioluminescence analysis of luciferase-expressing *E. coli* cells incubated with 100 μM CBT-OH and 100 μM D-Cystine for 30 min, followed by centrifugation and separation of supernatant and pellet at which point luminescence measurements were initiated and continued for 60 min. Non-centrifuged cells (CBT-OH + D-Cystine) were used as a control. (**d**) Luciferase-expressing *E. coli* cells treated with 100 μM CBT-OH and 100 μM D-Cystine in the presence and absence of IPTG induction, over 60 min. (**e**) Luciferase-expressing *E. coli* cells treated with 100 μM CBT-OH or 100 μM CBT-NH_2_ and 100 μM D-Cystine over 60 min. Cells treated with PBS and CBT only (CBT-OH and/or CBT-NH_2_) were used as controls, wherever appropriate. Data are represented as mean +/- SD (n = 3 independent samples in a single experiment).

Next, we set out to establish whether the luminescence signal generation was confined to within the cellular structure instead of extracellular recombination of CBT/D-Cys. Luciferase-expressing *E. coli* cells were first co-incubated with CBT-OH and D-Cystine for 30 min, after which, the cells were subjected to centrifugation, leading to the separation of the supernatant from the cellular pellet. The supernatant was then resuspended in phosphate-buffered saline (PBS). Luminescence measurements were then performed for an additional 60 min (**Fig. 2c**). Consistent with the expected cytosolic localization of the luciferase, luminescence signal detected in the supernatant was minimal, whereas the signal originating from the cellular pellet was significantly more pronounced. These observations are consistent with the intracellular nature of the signal and provide evidence that the luciferase enzyme remains localized within the cytoplasm throughout the entire duration of the assay.

We appreciated that the physicochemical properties of CBT could potentially be subjected to its own cellular accumulation barriers. We therefore sought to test two versions of CBT that have been described to be compatible with recombination with D-Cys to form luciferin. We evaluated CBT-OH and 6-amino-2-cyanobenzothiazole (CBT-NH_2_)^34^ as potential substrates for the click reaction (**Fig. 2b, e**). Notably, the *in vitro* rate constants for the reactions of the hydroxy- and amino-cyanobenzothiazole with L-Cys have previously been reported to be 3.2 and 2.6 M^-1^ s^-1^, respectively.^35^ Briefly, luciferase-expressing *E. coli* cells were co-incubated with the CBT variants and D-Cystine as described before. Our results showed that cells treated with CBT-NH_2_ produced a higher luminescence signal than those treated with CBT-OH (**Fig. 2d**). Interestingly, CBT-OH outperformed CBT-NH_2_ in an *in vitro* cell-free setup consistent with prior reports (**Fig. S2**). We pose that CBT-NH_2_ may have higher levels of accumulation relative to its hydroxy counterpart and it was therefore selected as the preferred CBT variants for subsequent experiments.

We then set out to empirically determine the optimum concentration of CBT-NH_2_ for DCCAA. Briefly, following IPTG induction, bacterial cells were washed and either co-incubated with varying concentrations of CBT-NH_2_ and 50 µM D-Cystine or incubated with CBT-NH_2_ alone. A concentration of 100 µM was considered to have a sufficient signal-to-noise ratio necessary for the assay and was selected for subsequent experiments (**Fig. 3a**). This concentration of CBT-NH_2_ was also found to not alter the cellular viability as determined by colony forming units (CFUs) analysis (**Fig. S3**). Next, a similar titration experiment was performed with D-Cystine using a constant level of CBT-NH_2_. Our results showed that both 50 µM and 100 µM of D-Cystine exhibited a favorable signal-to-noise ratio (**Fig. 3b**).

**Figure 3.**
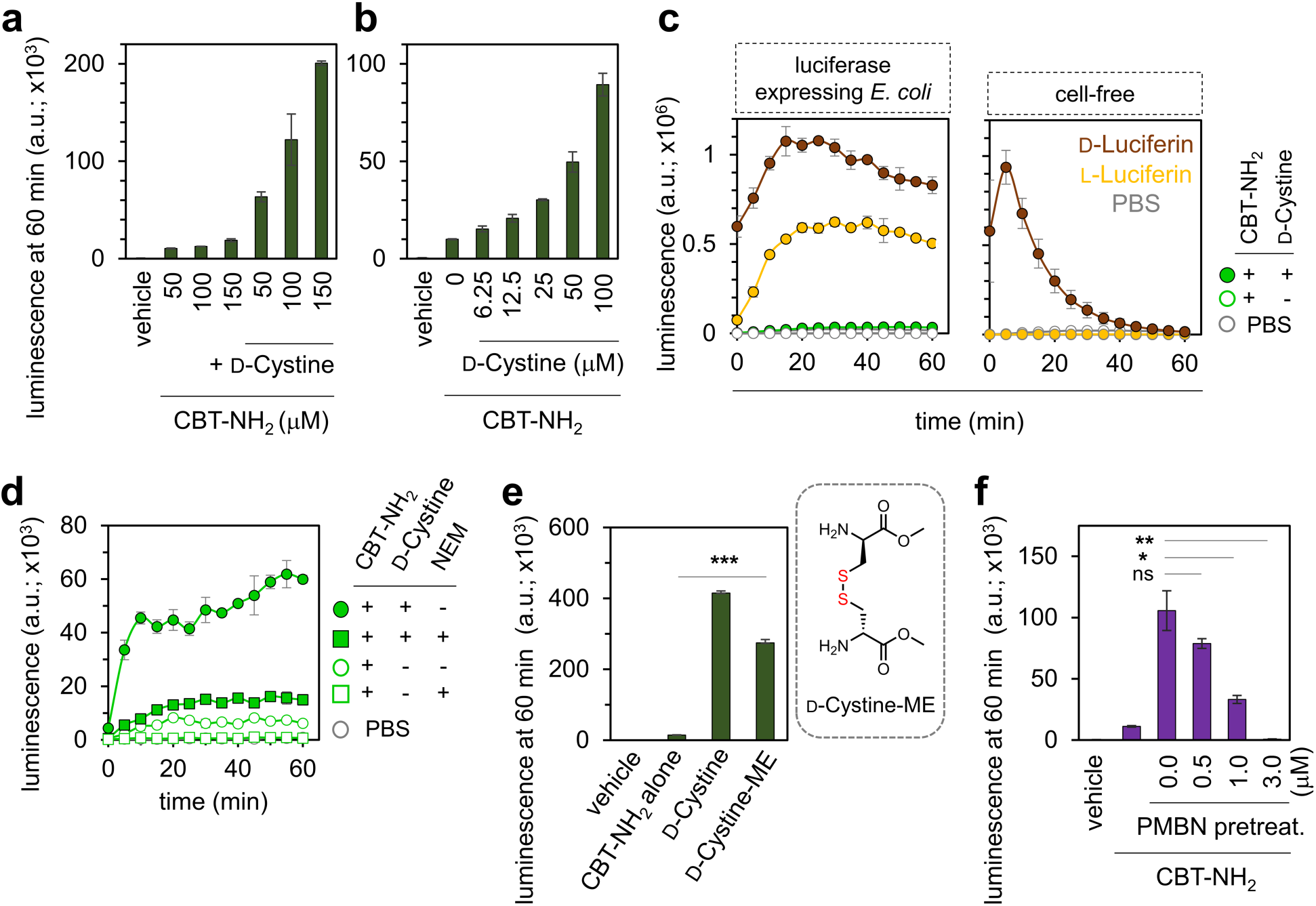
**(a)** Bioluminescence analysis of luciferase-expressing *E. coli* cells treated with 50 μM D-Cystine and varying concentrations of CBT-NH_2_ over 60 min at the 60 min time point. **(b)** Luciferase-expressing *E. coli* cells treated with 100 μM NH_2_-CBT and varying concentrations of D-Cystine at the 60 min time point. **(c)** Luciferase-expressing *E. coli* cells (left) or a cell-free assay with luciferase enzyme (right) treated with 10 μM L-Luciferin, 10 μM D-Luciferin or 10 μM D-Cystine with 100 μM CBT-NH_2_ (cell assay), over 60 min. **(d)** Luciferase-expressing *E. coli* cells treated with 100 μM D-Cystine and 100 μM CBT-NH_2_ or 100 μM CBT-NH_2_ only, in the presence and absence of a 30 min pretreatment with 100 μM NEM, over 60 min. **(e)** Luciferase-expressing *E. coli* cells treated with 100 μM D-Cystine or 100 μM D-Cystine-ME and 100 μM CBT-NH_2_ at the 60 min time point. **(f)** Luciferase-expressing *E. coli* cells treated with 100 μM D-Cystine and 100 μM CBT-NH_2_ in the absence and presence of a 30 min pretreatment with PMBN at different concentrations at the 60 min time point. Cells treated with PBS and CBT-NH_2_ only were used as controls, wherever appropriate. Data are represented as mean +/- SD (n = 3 independent samples in a single experiment). Statistical analysis performed by two-tailed t-test with Welch’s correction, * p ≤ 0.01, ** p ≤ 0.01, *** p ≤ 0.001, ns = not significant.

Through our initial assay development efforts, the background signal was higher than we had anticipated therefore we sought to investigate its potential source. We considered that it could be from intracellular pools of L-Cys combining with CBT-NH_2_ to form L-luciferin (**Fig. S4**). Previous reports have shown that L-luciferin can undergo epimerization in the presence of firefly luciferase and then act as a substrate for the enzyme.^36^ Indeed, upon co-incubation luciferase-expressing *E. coli* with L-luciferin, a signal lower than that of D-luciferin, yet significantly higher than the background, was observed (**Fig. 3c**). This observation suggests that L-luciferin undergoes epimerization within the cytoplasm of *E. coli*. In a cell free experiment, L-luciferin exhibited a baseline luminescence signal in the presence of purified luciferase, whereas D-luciferin exhibited a considerably elevated signal intensity (**Fig. 3c**). Notably, this observation is in agreement with the essential role of Coenzyme A (CoA) in the epimerization process of L-luciferin to D-luciferin catalyzed by luciferase.^37,38^

We next sought to evaluate the necessity for the D-Cys to be uncoupled prior to signal generation. At first, a cell free set up was evaluated by incubating D-Cystine with luciferase in the presence or absence of the reducing agent tris(2-carboxyethyl) phosphine (TCEP) (**Fig. S5**). The inclusion of TCEP was found to be essential for signal generation, providing evidence for the requirement of a reducing environment to promote signal production. A similar requirement for a reducing environment was next evaluated in cell.

For these experiments, *E. coli* was treated with N-ethylmaleimide (NEM), which covalently modifies cellular thiols and is expected to reduce the cellular pool of reducing agents including glutathione and L-Cys.^23,39^ Luciferase-expressing cells were pre-treated with NEM or PBS, followed by incubation with D-Cystine, as described previously. Pre-treatment with NEM resulted in a complete shutdown of signal (**Fig. 3d**). These results demonstrate that the reducing environment of the cell is required for signal generation in a manner that is consistent with the levels of cellular thiols. Moreover, we tested the effect of NEM on the background signal that emerged solely from the addition of CBT-NH_2_. Our findings demonstrate that NEM also effectively abolished the signal originating from CBT-NH_2_ (**Fig. 3d**), indicating that the presence of L-Cys most likely contributes to the observed background signal.

Considering the inherent characteristics of our initial test compound, D-Cystine, containing two carboxylic acid moieties, we explored the possibility of masking these groups to enhance accumulation. Masking negatively charged carboxylic acids through esterification is widely used as a permeability strategy to enhance the lipophilicity and passive membrane permeability of molecules, particularly, therapeutic agents with intracellular targets.^40,41^ Once inside the cell, the ester may be enzymatically hydrolyzed to the acid resulting in conversion to the parent compound. It is noteworthy that the introduction of a methyl group at the carboxylic position in D-luciferin, as seen in D-luciferin-methyl-ester, results in its failure to be recognized by the firefly luciferase enzyme.^42^ While the findings of Tippmann and colleagues^43^ suggest an absence of methyl esterases in *E. coli*, subsequent research by the Grimes laboratory^44^ utilizing methyl ester NAM derivatives in their peptidoglycan labeling approach, lends support to the presence of methyl esterases in *E. coli*.

To test the masking ability of methyl ester, we used DCCAA to compare D-Cystine and D-Cystine-methyl ester (D-Cystine-ME) (**Fig. S6**). Our results showed that cellular treatment with D-Cystine-ME led to signals well above background, suggestive of esterase unmasking of the methyl ester (**Fig. 3e**). Interestingly, the cellular signals were ca. 30% lower than that of unmasked D-Cystine. This could indicate that D-Cystine may be actively transported by an importer^45,46^ or the esterase processing is inherently slow. When the experiment was conducted in a cell-free system, the signal from D-Cystine-ME remained at the background level (**Fig. S7**). These findings provide support for the processing of ester groups in *E. coli* cells. The presence of esterases in *E. coli* bears relevance in the realm of drug development, especially in the potential utilization of a prodrug approach for antibiotics. Nevertheless, we believe that DCCAA can be generally leveraged to gain further insight into the substrate specificity and enzymatic activity of *E. coli* esterases.

The diderm cell envelope structure in Gram-negative bacteria, particularly the outer membrane, serves as a substantial barrier to the permeation of antibiotics with intracellular targets. Therefore, there is a critical need to identify and develop molecules that can disrupt this accumulation barrier. Such molecules can potentially be used as antibiotic adjuvants that can broadly improve activity. One example of molecule capable of perturbing the outer membrane is polymyxin B nonapeptide (PMBN), which is a modified form of Polymyxin B lacking the fatty acid tail. PMBN can permeabilize the outer membrane of *E. coli* at low, non-toxic concentrations.^47^ Moreover, it has also demonstrated the ability to enhance the efficacy of erythromycin and provide protection against Gram-negative bacteria in mice.^48^ We posed that the inclusion of PMBN in our assay would increase the permeability of bacterial cells to D-Cystine, leading to an amplified signal response.

To test the potential impact of PMBN, luciferase expressing *E. coli* cells were pre-treated with PMBN at increasing concentrations and the assay was carried out as described previously (**Fig. 3f**). Unexpectedly, a dose-dependent decrease in luminescence signal with increasing concentrations of PMBN was observed. This trend was also observed when cells were, instead, treated with D-luciferin (**Fig. S8**). Crucially, we noted no loss of cellular viability with the same concentrations of PMBN (**Fig. S9**). We wondered whether PMBN could be leading to the release of components critical to DCCAA. We observed a dose-dependent increase in outer membrane permeation as measured by nitrocefin (**Fig. S10**). A similar pattern was observed in the case of a SYTOX Green assay, which is a high-affinity nucleic acid stain typically used to assess membrane integrity (**Fig. S11**). Notably, treatment of *E. coli* cells with PMBN has been previously reported to cause the release of intracellular, low-molecular weight substances such as free amino acids and uracil.^49^ Hence, we hypothesize that the observed reduction in signal may be due to the leakage of substrates, specifically CBT or D-Cystine (or D-luciferin) from the cells. Additionally, it could also stem from the release of ATP from the cells which is integral for the oxidation of D-luciferin by luciferase.^50,51^

Having optimized the assay parameters of DCCAA and shown its ability to respond to the model molecule D-Cystine, we next set out to leverage this assay to evaluate the accumulation of structural motifs that are relevant for clinical applications. For this, a small panel of antibiotics (including ciprofloxacin, puromycin, linezolid, and rifamycin B) were synthesized and tagged with a disulfide-linked D-Cys (**Fig. 4a**). In all, this panel included a range of molecules that spanned variable physicochemical properties. Before performing the cellular assay, we aimed to test the release of D-Cys in a cell-free experiment. It is noteworthy that, in the absence of a permeability barrier, when exposed to a reducing agent, all four compounds were expected to generate an identical luminescence signal at the same concentration. This should be because of the release of an equal number of D-Cys molecules from each compound.

**Figure 4.**
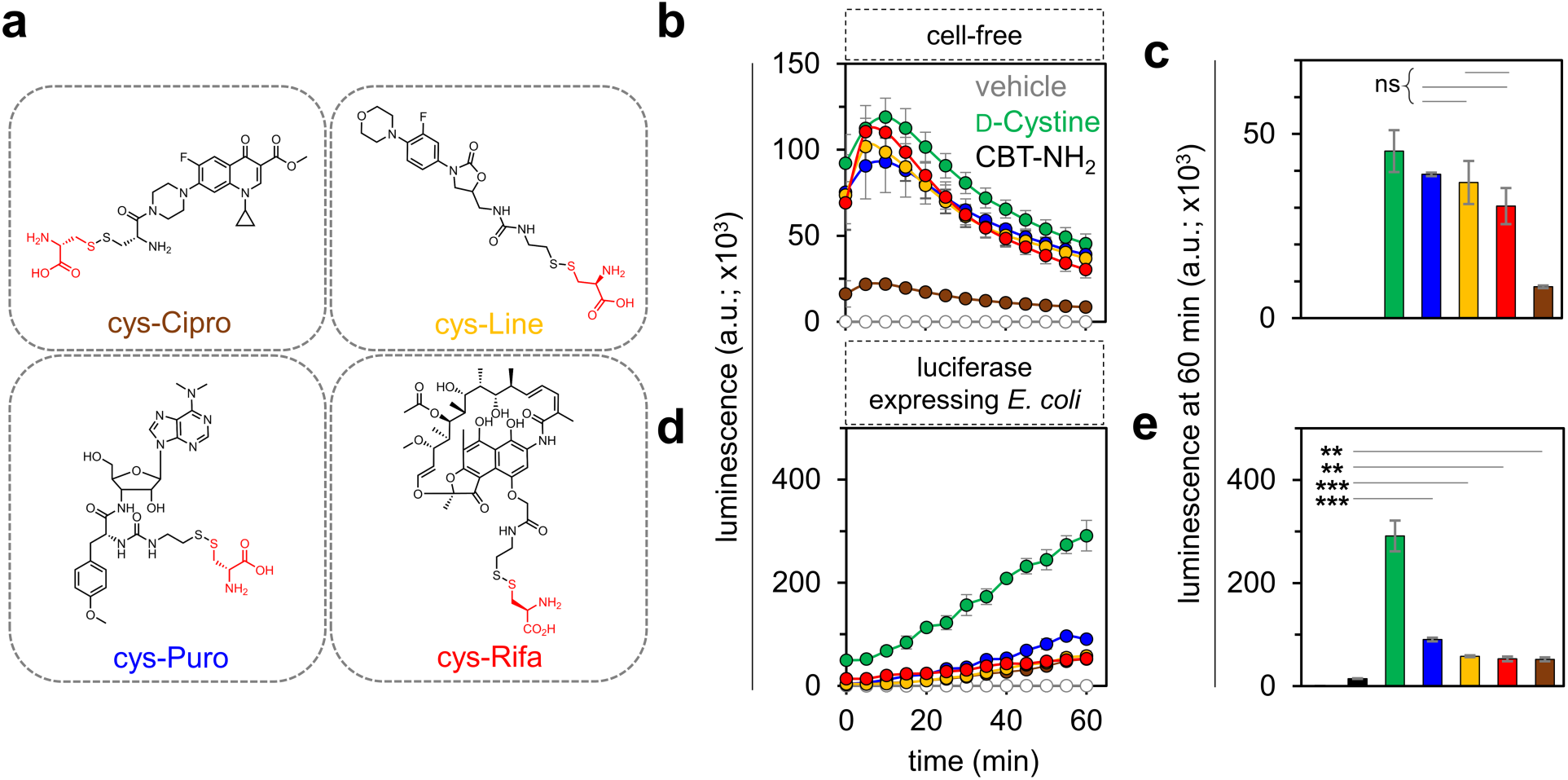
**(a)** Structures of the antibiotic conjugates tagged with a D-Cys moiety via a disulfide linkage. **(b)** Bioluminescence analysis of a cell-free assay with luciferase enzyme treated with 25 μM D-Cystine or 25 μM antibiotic conjugates and 25 μM CBT-NH_2_ over 60 min and **(c)** at the 60 min time point upon treatment with 1mM TCEP. **(d)** Bioluminescence analysis of luciferase-expressing *E. coli* cells treated with 50 μM D-Cystine or 50 μM of antibiotic conjugates with 100 μM CBT-NH_2_ over 60 min and **(e)** at the 60 min time point. Cells treated with PBS and CBT only (CBT-NH_2_) were used as controls, wherever appropriate. Data are represented as mean +/- SD (n = 3 independent samples in a single experiment). Statistical analysis performed by two-tailed t-test with Welch’s correction, ** p ≤ 0.01, *** p ≤ 0.001, ns = not significant.

Interestingly, our cell-free results revealed an unexpected result (**Fig. 4b**). While there was no significant difference in luminescence signal for puromycin, rifamycin B, and linezolid conjugates, the signal for ciprofloxacin was much lower (**Fig. 4c**). We hypothesized that the residual covalently tagged D-Cys in the ciprofloxacin conjugate after the breakage of the disulfide bond, could potentially undergo a non-productive click reaction with CBT-NH_2_ (**Fig. S12**) thereby reducing the effective concentration of CBT. Nonetheless, all four antibiotics were evaluated in the cellular assay. As before, the evaluation of these molecules was conducted by co-incubating luciferase-expressing bacterial cells with CBT-NH_2_ and the antibiotic conjugates, and subsequent luminescence measurements were made. It is noteworthy that all four conjugates demonstrated noticeable permeation exceeding the baseline signal as seen in **Fig. 4d**, as indicated by the luminescence signals observed at the 60 min time point (**Fig. 4e**). Notably, addition of the antibiotic conjugates to the luciferase expressing *E. coli* cells was found to be minimally disruptive to their cell envelope structure, as evidenced by a SYTOX Green assay (**Fig. S13**).

We then proceeded to develop a kinetic model for molecular uptake of the conjugated antibiotics. The model serves as the foundation for a nonlinear least-squares analysis of our data. Specifically, the model accounts for the accumulation of each molecule in the cytoplasm of *E. coli* and the subsequent click and luciferase-based bioluminescence enzymatic reactions. Of significance, the time-dependent luminescence response (I*_Lum._*) can be characterized by Eq. (1), where ε_0_ and I_0_ respectively represent the luminescence coefficient factor and baseline signal in our experiments. D-Luciferin ^(^*^)^ denotes the oxidized D-luciferin molecule which is associated with the luminescence signal.

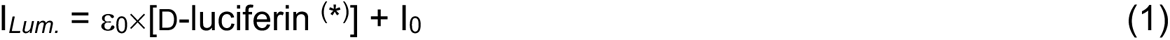

Time-dependent luminescence signals are then utilized to derive a solution with physical significance for the specified differential equations, as detailed in the SI. Special attention is given to estimating the accumulation rate constant, denoted as k_acc_ (min^-1^), and the reduction rate constant, denoted as k_red_ (min^-1^). Values for the click reaction rate constant, k_click_ (156 M^-1^min^-1^)^35^ and the enzymatic reaction rate constant, k_enz_ (96 min^-1^)^52^ were adopted from the literature.

To illustrate how k_acc_ and k_red_ affect time-dependent luminescence signals, we selectively adjusted the associated transport rates in a set of simulated kinetic responses. In **Fig. 5a**, a sequence of simulated responses is presented, incorporating various k_acc_ values ranging from 0.01 to 1.00 min^-1^, paired with each k_red_ value set at 0.01, 0.10, and 0.50 min^-1^. Importantly, an offset of 3 min was introduced in the experimental data to account for the time delay between the beginning of the assay and the first luminescence reading, owing to logistics of pipetting, mixing, etc. Additionally, all signals, experimental and simulated, were normalized with respect to their values at 63 min for the purpose of this model. As depicted, irrespective of the k_red_ values, the curves exhibit exponential behavior at lower k_acc_ values, transitioning gradually into a semi-linear shape at higher k_acc_ values.

**Figure 5.**
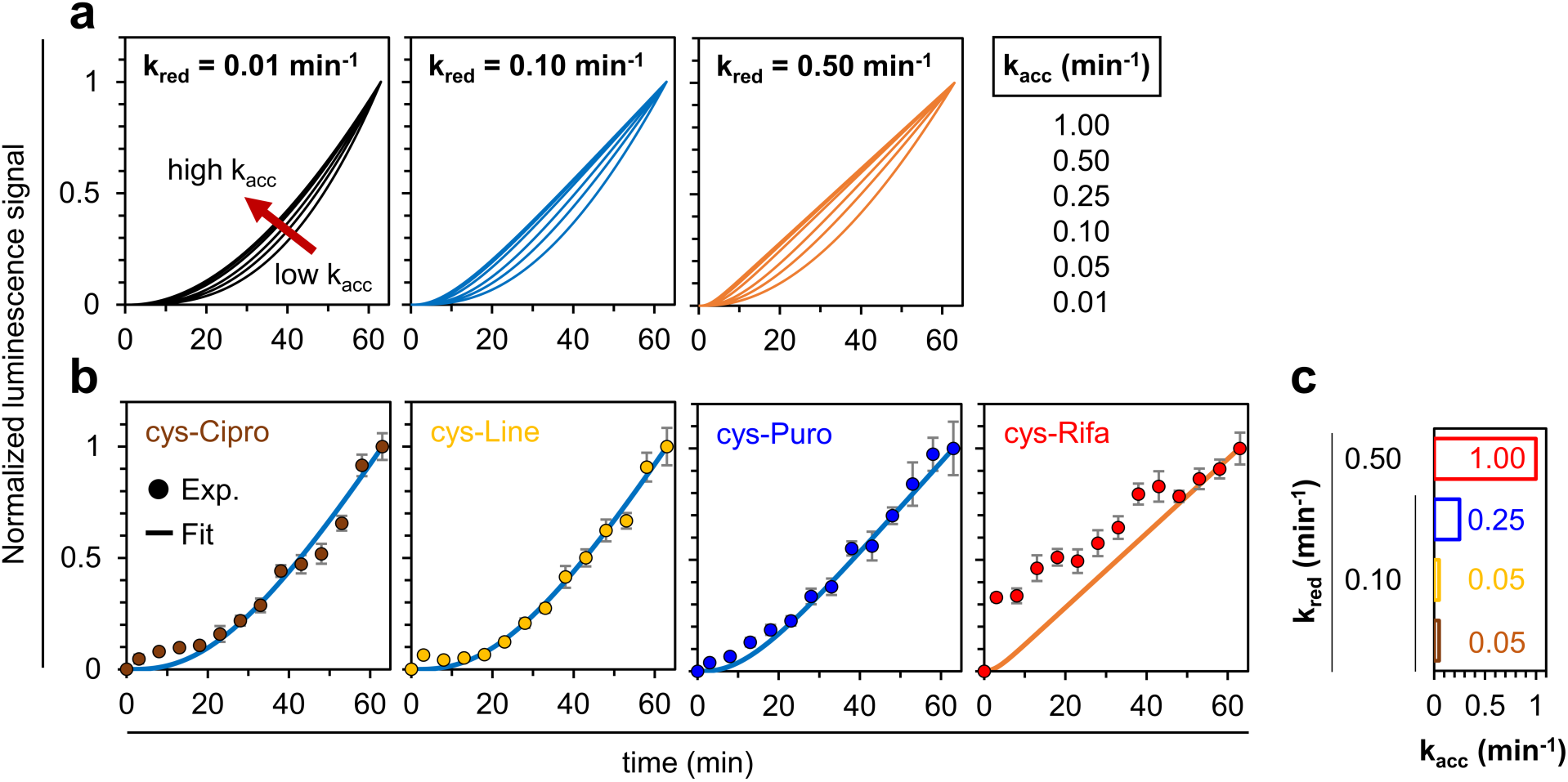
**(a)** Simulated luminescence traces, based upon the model for molecular accumulation in bacteria, with gradually increasing rate constants k_acc_ and k_red_. **(b)** Experimental time-resolved luminescence traces illustrating the accumulation of test molecules in *E. coli*. The best-fitted lines from simulations are overlaid. The corresponding values for k_acc_ and k_red_ for each, are depicted in **(c)**.

Next, we sought optimal fits for each time-resolved signal, representing the averages from four independent trials. In **Fig. 5b**, the most accurate fitting curves are presented, superimposed on each signal. Notably, for the ciprofloxacin-methyl-ester, linezolid, and puromycin conjugates, the best-estimated k_acc_ values were respectively determined to be 0.05, 0.05, and 0.25 min^-1^, each accompanied by a k_red_ of 0.10 min^-1^ (**Fig. 5c**). These findings imply that the membrane transport rates for conjugated ciprofloxacin methyl ester and linezolid are five times lower than that of conjugated puromycin. Significantly, despite linezolid being known for its vulnerability to efflux pump activity, we observed its accumulation behavior in our assay. This may be attributed to the possibility that our luminescence readout process operates on a faster timescale than its efflux out of the cell or that efflux pumps do not effectively recognize linezolid when tagged with a D-Cys moiety. As far as D-Cys is concerned, as expected, it displayed a higher k_acc_ value of 1.00 min^-1^ and k_red_ value of 0.50 min^-1^, suggesting a faster accumulation and reduction processes, compared to the conjugated antibiotics (**Fig. S14**). Unexpectedly, the rifamycin B, when compared to the other three components, displayed slightly distinct kinetics. The best-fitted curve revealed a rate constant similar to that of D-cystine (i.e., k_acc_ = 1.00 min^-1^; k_red_ = 0.50 min^-1^). This may be due to a difference in its membrane transport mechanism or as a result of the property of the molecule itself, given its larger size. Significantly, molecules with comparable molecular weights to our antibiotic conjugates, such as malachite green (329.46 Da), have been documented to exhibit a transport rate of approximately 4.2 min^-1^ and 0.013 min^-1^ through the outer- and inner-membranes of *E. coli*, respectively.^53^ In contrast, a similar molecule without a permanent dipole moment, namely crystal violet (372.54 Da; net charge of 1+), displays inner-membrane transport rates that can be orders of magnitude smaller.^54^

## Conclusion

Quantifying intracellular compound accumulation within the bacterial cytosol especially during the development phase of antibiotics with cytosolic targets is crucial for the development of efficacious drugs with usable potency against their targets. Here, we present DCCAA, a luminescence-based method enabling real-time monitoring of antibiotic intracellular accumulation in live Gram-negative bacterial cells. In this technique, the molecule of interest is conjugated with a D-Cys moiety through a disulfide bond, susceptible to reduction within the cytosolic milieu. The liberated D-Cys then engages in a click reaction with CBT, co-administered with the tagged molecule, resulting in the generation of D-luciferin. This substrate undergoes oxidation by luciferase, emitting light in the process, as a measurable indicator of intracellular antibiotic accumulation dynamics (**Fig. 1a**). Additionally, our results showed compelling evidence supporting the presence of methyl esterases in *E. coli*. This investigation highlights the potential use of DCCAA to decipher the presence and/or the substrate-specificity of esterases in different bacterial species. We were also able to elucidate the membrane disruption activity of an antimicrobial compound. Specifically, we investigated the activity of PMBN, a known Gram-negative outer membrane disrupting agent, via DCCAA. Lastly, through DCCAA we were able to monitor the intracellular accumulation of the D-Cys conjugates of ciprofloxacin methyl ester, linezolid, puromycin and rifamycin B. We found all four antibiotic conjugates tested to display a signal above background, indicating accumulation in the cytosol of bacteria. Through kinetic analysis, we found that the puromycin conjugate displayed a higher transport rate than the linezolid and ciprofloxacin-methyl-ester conjugates while the rifamycin B conjugate displayed a distinct kinetic profile, unlike the other compounds, likely owing to a distinct membrane transport process or its larger size.

In conclusion, we propose that DCCAA can serve not only to elucidate the real-time intracellular accumulation dynamics of compounds tagged with a D-Cys in live bacterial cells but also to potentially unveil the membrane disruption abilities of untagged compounds. Furthermore, DCCAA can also be employed to reveal the presence and substrate specificity of esterase activity in live bacterial cells.

## Acknowledgement

This study was supported by the NIH grant GM124893-01 (M.M.P.).

## Supporting Information

Additional figures, tables, and materials/methods are included in the supporting information file.

