## Supporting Information for "Bioluminescence-Based Determination of Cytosolic Accumulation of Antibiotics in *Escherichia coli*"

### Table of Contents


| | |
|---|---|
| **Supporting Figures** | **S3-S16** |
| **Figure S1:** CBT and D-Cys reaction to form D-luciferin and subsequent conversion to Oxyluciferin by firefly luciferase, emitting light | **S3** |
| **Figure S2:** $NH_2$-CBT vs OH-CBT in a cell-free luciferase assay | **S4** |
| **Figure S3:** CFU analysis of cells treated with $NH_2$-CBT | **S5** |
| **Figure S4:** CBT and L-Cys reaction, L-luciferin to D-luciferin conversion by firefly luciferase | **S6** |
| **Figure S5:** Effect of TCEP in a cell-free luciferase assay with $NH_2$-CBT and D-Cystine | **S7** |
| **Figure S6:** Conversion of D-Cystine-ME to D-Cys followed by conversion to D-luciferin upon entry into the *E. coli* cytosol | **S8** |
| **Figure S7:** Cell-free luciferase assay with $NH_2$-CBT and D-Cys or D-Cys-Methyl-Ester | **S9** |
| **Figure S8:** Effect of PMBN pretreatment on luciferase-expressing *E. coli* cells treated with D-luciferin | **S10** |
| **Figure S9:** CFU analysis of cells treated with PMBN | **S11** |
| **Figure S10:** Nitrocefin assay for PMBN impact on *E. coli* outer membrane integrity | **S12** |
| **Figure S11:** SYTOX-Green assay for PMBN impact on *E. coli* cell-envelope integrity | **S13** |
| **Figure S12:** Non-productive click reaction of residual D-Cys in the ciprofloxacin conjugate (cys-Cipro) with CBT-$NH_2$ post-disulfide bond cleavage | **S14** |
| **Figure S13:** SYTOX-Green assay for antibiotic conjugate impact on *E. coli* cell-envelope integrity | **S15** |
| **Figure S14:** Simulation fit for the accumulation of D-Cystine in *E. coli* | **S16** |
| **Methods** | **S17-S19** |
| Materials | **S17** |
| Transformation of FLUC2 pET28a into *E. coli* | **S17** |
| Luciferase Protein Expression in *E. coli* | **S17** |
| Evaluation of Luciferase Protein Expression in *E. coli* via SDS-PAGE | **S17** |

| | |
|---|---|
| Bioluminescence-Based Permeability Assays in Luciferase Expressing *E. coli* | **S18** |
| Cell-free Luciferase Assays | **S18** |
| CFU Analysis | **S18** |
| Nitrocefin Assay | **S19** |
| SYTOX-Green Assay | **S19** |
| Kinetic Model of Cytosolic Accumulation of Small Molecules in *E. coli* | **S19** |
| **Synthesis and Characterization** | **S20-S34** |
| General Methods | **S20** |
| cys-Cipro | **S21** |
| cys-Line | **S27** |
| cys-Puro | **S31** |
| cys-Rifa | **S34** |
| **References** | **S38** |

### Supporting Figures

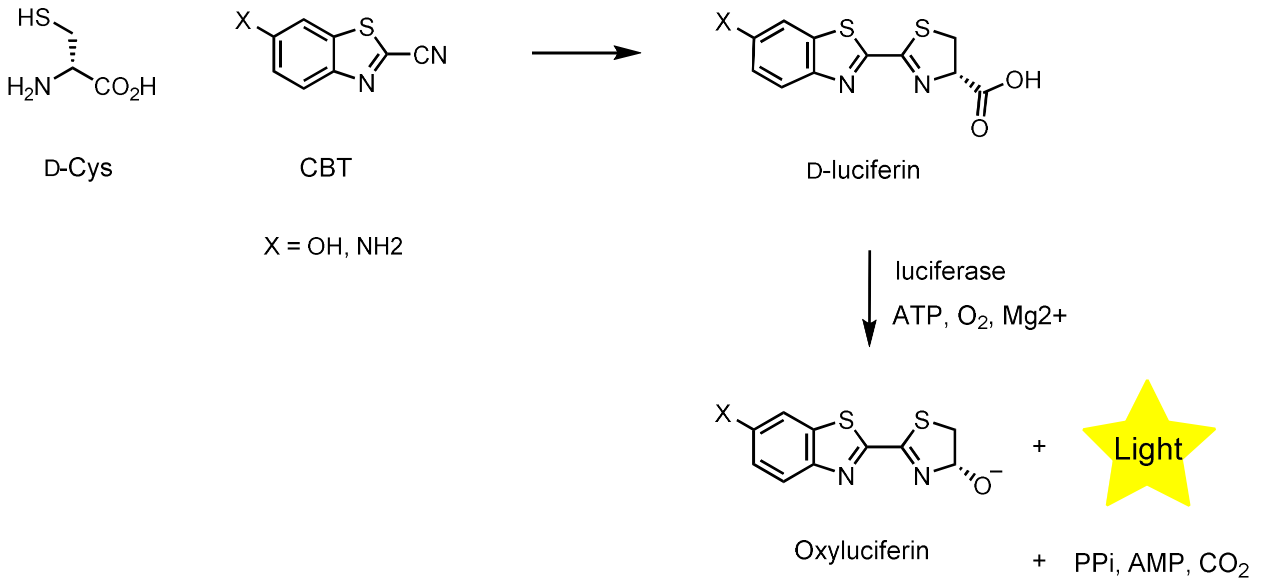


**Figure S1.** Schematic depicting the reaction between CBT and D-Cys followed by the enzymatic conversion of D-luciferin to oxyluciferin by firefly luciferase, resulting in the emission of light.

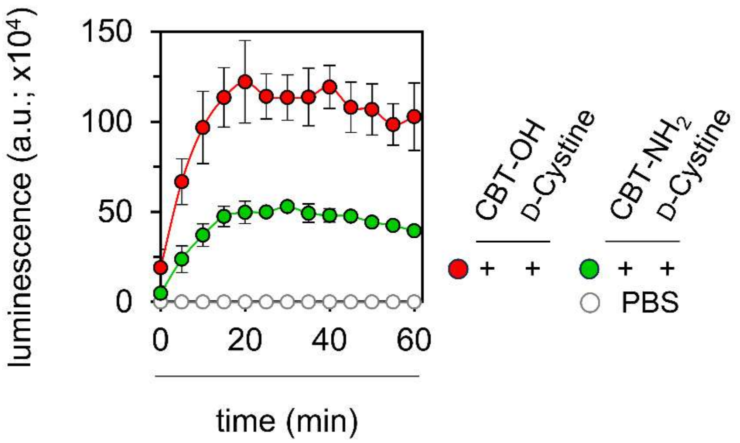


**Figure S2.** Bioluminescence analysis of a cell-free assay with luciferase enzyme treated with 25 μM $NH_2$-CBT or OH-CBT and 25 μM D-Cystine, over 60 minutes. Treatment with PBS was used as the blank control. Data are represented as mean +/- SD (n = 3 independent samples in a single experiment).

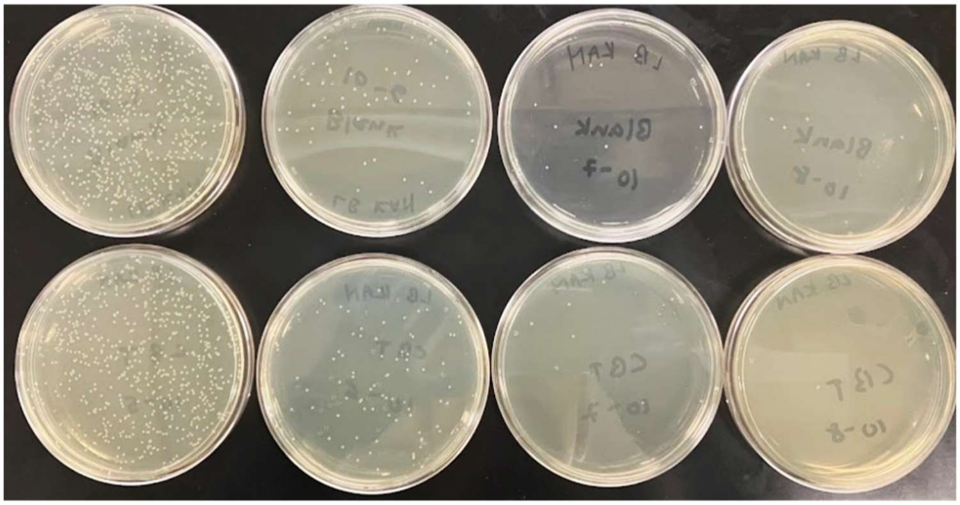

**Figure S3.** E. coli cells were induced with IPTG and were untreated (top row) or treated with 100 μM $NH_2$-CBT (bottom row) for 1 h at 37 °C in PBS. Next, serial dilutions of the cultures were made, and cells were spread on agar/LB plates with kanamycin and placed in a 37 ºC incubator for 15 hours. Images of four sets of treated and untreated agar plates are shown. At $10^{-6}$ dilution, the CFU counts were 83 for $NH_2$-CBT-treated and 59 for untreated.

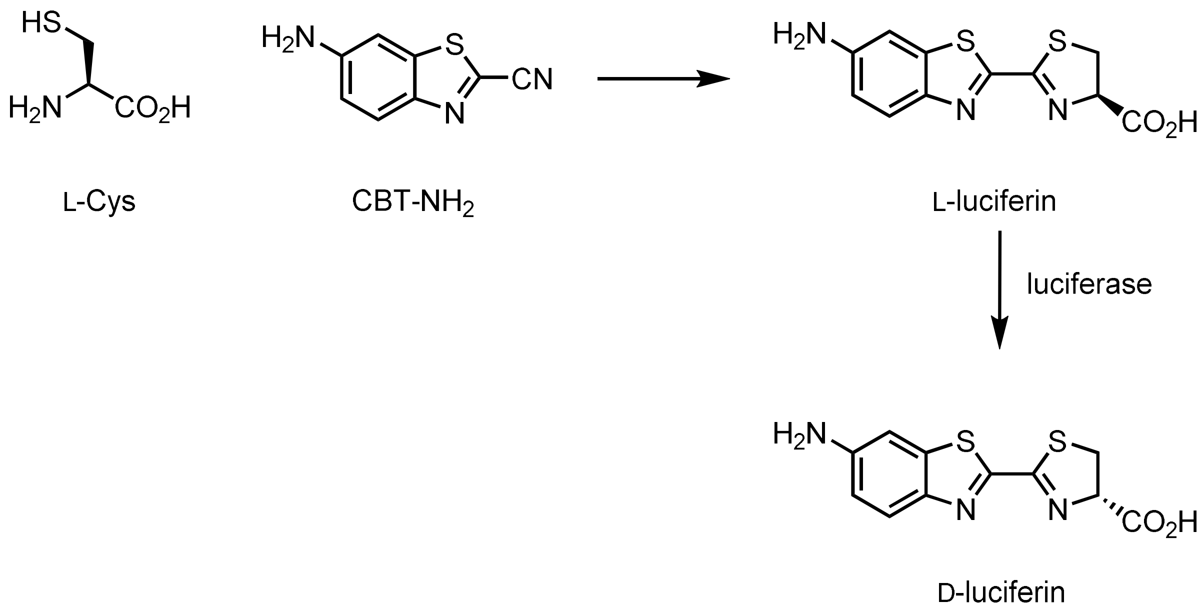


**Figure S4.** Schematic depicting the reaction between CBT and L-Cys followed by the enzymatic conversion of L-luciferin to D-luciferin by firefly luciferase.

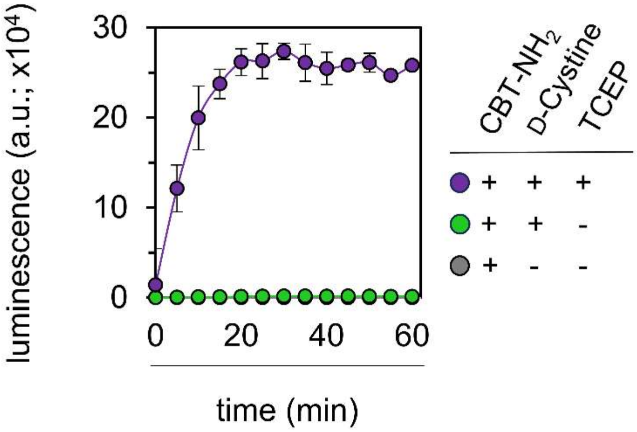


**Figure S5.** Bioluminescence analysis of a cell-free assay with luciferase enzyme treated with 100 μM $NH_2$-CBT and 100 μM D-Cystine in the presence and absence of 1mM TCEP over 60 minutes. Data are represented as mean +/- SD (n = 3 independent samples in a single experiment).

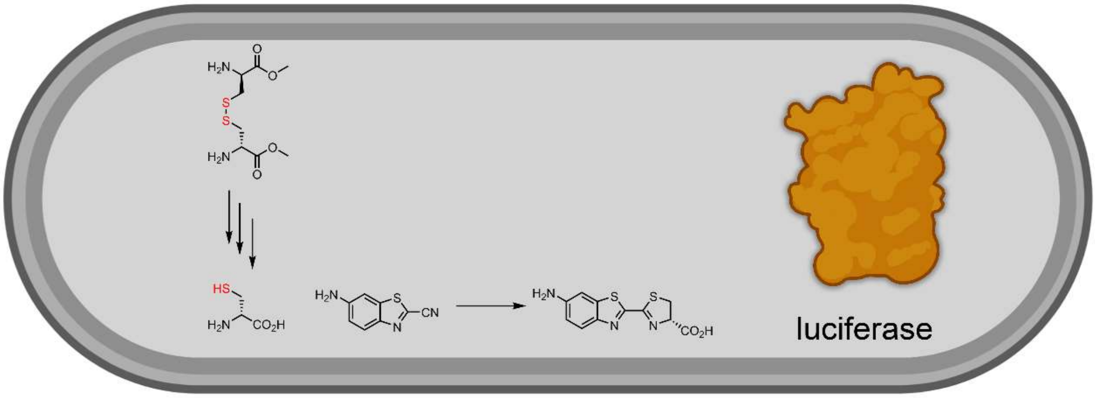


**Figure S6.** Conversion of D-Cystine-ME to D-Cys upon entry into the *E. coli* cell expressing luciferase and subsequent reaction of D-Cys with CBT to form D-luciferin.

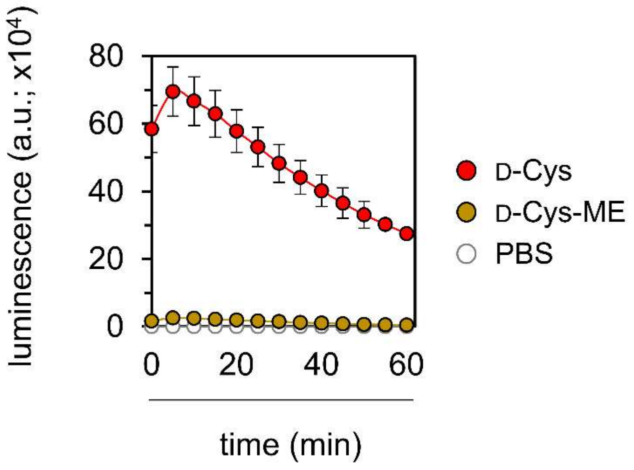


**Figure S7.** Bioluminescence analysis of a cell-free assay with luciferase enzyme treated with 25 µM $NH_2$-CBT and 25 µM D-Cys or D-Cys-ME over 60 minutes. Data are represented as mean +/- SD (n = 3 independent samples in a single experiment).

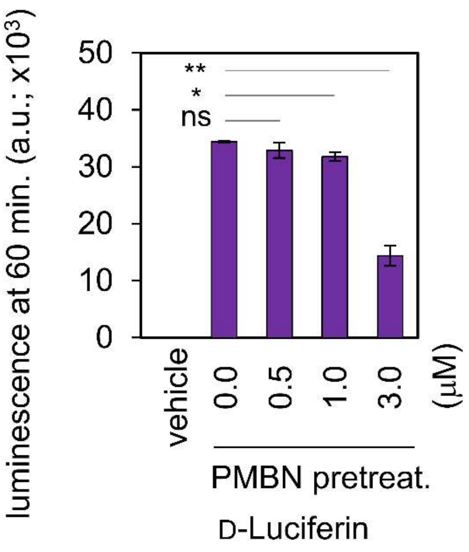


**Figure S8.** Bioluminescence analysis of luciferase-expressing *E. coli* cells treated with 10 μM D-luciferin in the absence and presence of a 30-minute pretreatment with PMBN at the 60-minute time point. Cells treated with PBS (vehicle) and CBT only ($NH_2$-CBT) were used as controls. Data are represented as mean +/- SD (n = 3 independent samples in a single experiment). Statistical analysis by two-tailed t-test with Welch's correction, * $p \leq 0.01$, ** $p \leq 0.01$, ns = not significant.

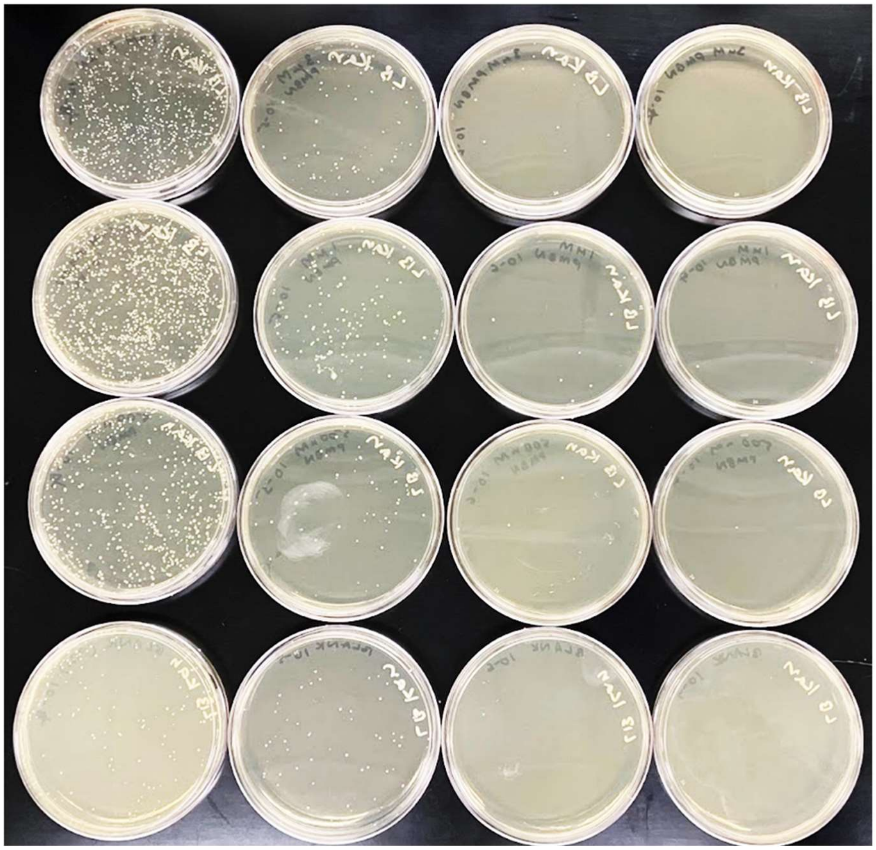

**Figure S9.** E. coli cells were induced with IPTG and were untreated (first, i.e., bottom row) or treated with 0.5 μM (second row), 1 μM (third row), and 3 μM (fourth row) PMBN for 30 minutes at 37 °C in PBS. Next, serial dilutions of bacterial culture were made, and cells were spread on agar/LB plates with kanamycin and placed in a 37 ºC incubator for 15 hours. Images of four sets of treated and untreated agar plates are shown. At $10^{-6}$ dilution, the CFU counts were 62 for untreated, 66 for 0.5 μM, 107 for 1 μM, and 80 for 3 μM.

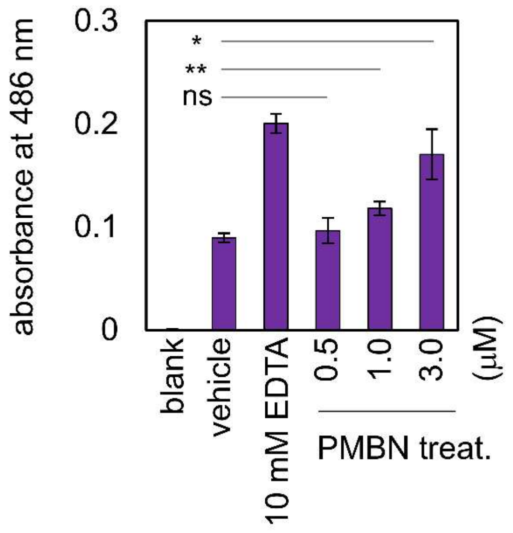


**Figure S10.** Luciferase-expressing E. coli cells were co-treated with 50 μg/mL nitrocefin and PMBN at different concentrations for 30 minutes. Cells treated with 50 μg/mL nitrocefin and PBS or 10 mM EDTA were used as the negative (vehicle) or positive control, respectively. Untreated cells were used as the blank. Data are represented as mean +/- SD (n = 3 independent samples in a single experiment). Statistical analysis by two-tailed t-test with Welch's correction, * $p \leq 0.01$, ** $p \leq 0.01$, ns = not significant.

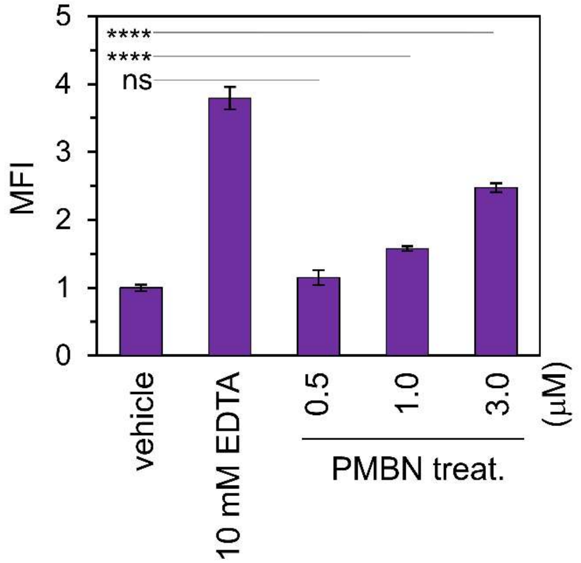


**Figure S11.** Luciferase-expressing E. coli cells were treated with SYTOX Green in the presence of a 30-minute pretreatment with PMBN at different concentrations. Cells treated with PBS or 10 mM EDTA prior to treatment with SYTOX Green were used as the negative or positive control, respectively. Data are represented as mean +/- SD (n = 3 independent samples in a single experiment). Statistical analysis by two-tailed t-test with Welch's correction, **** $p \leq 0.0001$, ns = not significant.

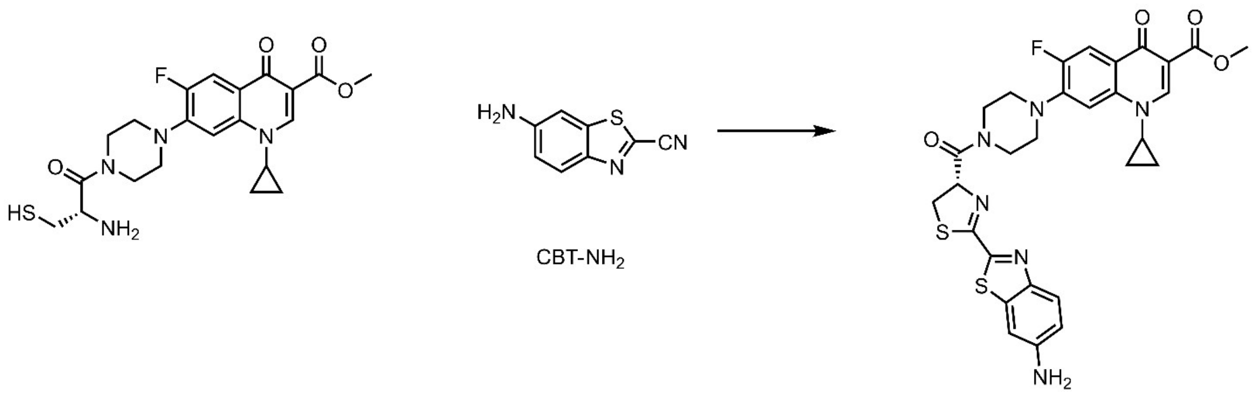


**Figure S12.** Post-disulfide bond cleavage, residual covalently tagged D-Cys in the ciprofloxacin conjugate (cys-Cipro) may engage in a non-productive click reaction with CBT-$NH_2$.

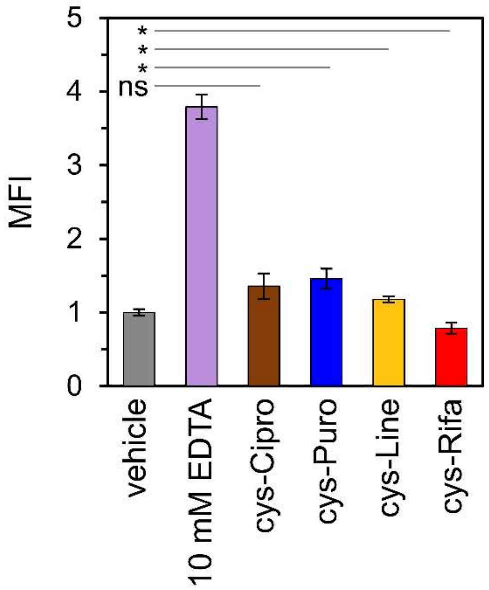


**Figure S13.** Luciferase-expressing E. coli cells were treated with SYTOX Green in the presence of a 1-hour pretreatment with the antibiotic conjugates at 50 μM. Cells treated with PBS or 10 mM EDTA prior to treatment with SYTOX Green were used as the negative or positive control, respectively. Data are represented as mean +/- SD (n = 3 independent samples in a single experiment). Statistical analysis by two-tailed t-test with Welch's correction, * $p \leq 0.01$, ns = not significant.

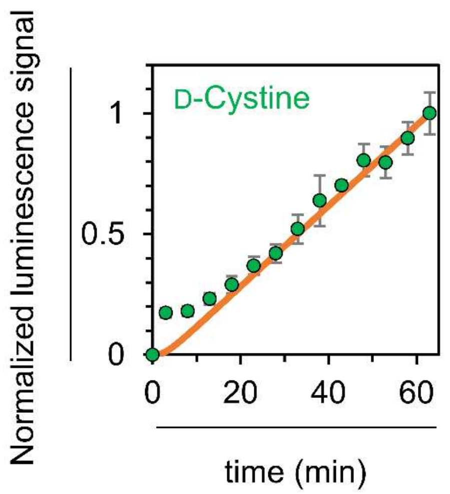


**Figure S14**. Experimental time-resolved luminescence trace illustrating the accumulation of D-Cystine in *E. coli*. The best-fitted line from simulations corresponding to a $k_{inter}$ value of 1.00 $min^{-1}$ and a $k_{red}$ value of 0.50 $min^{-1}$ is overlaid.

### Methods

**Materials.** The FLUC2 pET28a expression plasmid was obtained from the Ai lab at the University of Virginia. All reagents for assays, bacterial growth and chemical building blocks for synthesis were purchased either from ChemImpex, TCI Chemicals, ThermoFisher, ApexBio or Sigma-Aldrich. Luciferase Enzyme (Catalog No. E1701) was purchased from Promega. CBT-$NH_2$ (Catalog No. sc-233526) was purchased from ChemCruz. CBT-OH was purchased from Acros Organics (Catalog No. 450940010). D-luciferin (Catalog No. LUCK-250) and L-luciferin (Catalog No. L-127-10) were purchased from GoldBio. Deuterated solvents were used as received from Cambridge Isotopes.

**Transformation of FLUC2 pET28a into *E. coli*.** For expression of the luciferase protein in E. coli, the FLUC2 pET28a plasmid was first transformed into DH5α *E. coli* cells for amplification. In short, 50 µL of competent DH5α *E. coli* cells and 2-5 µL plasmid were added to an Eppendorf tube and kept on ice. After 30 minutes, a water bath was heated to 42 °C and the tube containing the cells and plasmid was placed in the bath for 30 seconds followed by another 2 minutes on ice. One mL of sterile LB medium was added to the tube, mixed, and transferred to a culture tube which was incubated at 37°C for 1 hour. Subsequently, 25-200 µL of cells from this tube were grown on LB/agar plates with kanamycin (50 µg/mL) at 37 °C overnight. Individual colonies were then picked from these plates and grown overnight in LB broth with 50 µg/mL kanamycin at 37°C. The following day, a ZymoPURE Plasmid Miniprep Kit was used to extract the plasmid from the DH5α *E. coli* cells. A transformation was then performed as above with competent BL21(DE3) *E. coli* cells for optimal expression. Glycerol stocks of both BL21 and DH5α *E. coli* cells were prepared by mixing 1 mL of overnight growth with 1 mL of 60% glycerol in water.

**Luciferase Protein Expression in *E. coli*.** Sterile culture tubes each containing 3 mL of LB medium, and 50 µg/mL kanamycin were inoculated with a stab of the transformed BL21 luciferase-expressing *E. coli* glycerol stock and incubated at 37 °C overnight. The next morning, the cells were diluted at a 1:10 ratio into fresh LB broth containing 50 µg/mL kanamycin and were grown at 37 °C for 3 h, or until the optical density at 600 nm reached a value between 0.6-0.8. Cultures were then induced with 1 mM isopropyl β-d-1-thiogalactopyranoside (IPTG) at 37 °C for 2 h in a shaker incubator to induce protein expression.

**Evaluation of Luciferase Protein Expression via SDS-PAGE.** Overnight cultures luciferase-expressing *E. coli* were grown and diluted the next morning as described above. When the diluted cells reached an OD600 between 0.6-0.8, IPTG was added to one culture tube to induce protein expression. Both plus and minus IPTG cultures were then incubated at 37 °C for 2 or 26 hours. On the same day that these cells were induced, another two overnight cultures were grown. The next day, these cells were diluted, and IPTG was used to induce protein expression of one culture for two hours at 37 °C. 1 mL of media was collected from the plus and minus IPTG samples from both time points. These were centrifuged at 3300g for 3 minutes in a HERAEUS Multicentrifuge X1 centrifuge (Thermo Fisher Scientific). The supernatant was discarded, and the cells were resuspended in 400 µL 1× phosphate buffered saline (PBS) at pH 7.2. 40 µL of the cell resuspension was mixed with 10 µL of a 5X SDS-PAGE sample loading buffer. This mixture was then boiled for 5 minutes to denature the proteins. To run the gel, 8 µL of Thermo Scientific™ PageRuler™ Plus Prestained Protein Ladder, 10 to 250 kDa, was added to the first well. The samples were loaded at 20 µL, and then gel was run at 240 V for 30 minutes.

**Bioluminescence-Based Permeability Assays in Luciferase Expressing *E. coli*.** Overnight cultures luciferase-expressing *E. coli* were grown, and luciferase protein expression was induced as above. Subsequently, washing was performed by first pelleting the cells in the HERAEUS Multicentrifuge X1 centrifuge (Thermo Fisher Scientific) at 3300g for 3 minutes. The supernatant was then discarded, and the cells were resuspended in 1× PBS (volume same as original culture). This was repeated 2 times. The cells were resuspended after the last wash in 1× PBS. In NEM/PMBN pretreatment experiments, a 30-minute incubation with the appropriate compound was performed at this point at the desired concentrations (100 µM for NEM; 0.5, 1 and 3 µM for PMBN). Cells were then washed and resuspended in 1× PBS in the same manner as described above. In a 96-well, black, flat-bottomed plate, the molecules of interest were then added to obtain the final desired concentration in a total volume of 100 µL. Note that stock solutions of molecules of interest were prepared in 1× PBS. 1× PBS was also added to any blank and negative control wells, to ensure the same total volume and number of cells in all the wells. A stock solution of CBT (6-Amino-2-cyanobenzothiazole) was prepared in N, N-Dimethylformamide (DMF), and appropriate volumes were added to the culture tubes containing cells to obtain the desired concentrations. Then, the same volume of cells was pipetted into each well, mixed, and placed in the BioTek Synergy H1 Microplate Reader. The instrument was set to run on the endpoint/kinetic luminescence setting. Luminescence was set to read every 5 minutes for the run, usually for 60 minutes. For the assay to test the luminescence of the pellet and supernatant of centrifuged *E. coli* cells, the plate was read for 30 minutes following which a set of wells treated with CBT and D-Cystine were moved to another plate and pelleted as described above. The supernatant was then carefully separated and re-added to the plate. The pellet was resuspended and also re-added to the plate. Non-pelleted cells treated with CBT and D-Cystine served as one of the controls. Luminescence in this case was then read for another 60 minutes. For all luminescence readings via the BioTek Synergy H1 Microplate Reader, luminescence fiber was the optics type. The gain was set to 240, and the integration time was set to 0:01:00. The temperature was set to 37 °C.

**Cell-free Luciferase Assays**. The 14.9 mg/mL luciferase enzyme stock received from Promega was diluted in Tris buffer (pH 8) to prepare 0.4 mg/mL aliquots (40µL each) which were then stored in -80ºC. All assays were performed in Tris buffer (pH 8) in 96-well, black, flat-bottomed plates. Molecules of interest were added to obtain the final desired concentration in a total volume of 100 µL. Appropriate amounts of $MgSO_4$ (100 mM), freshly made ATP (20 mM) and Luciferase stocks (0.4 mg/mL**)** prepared in Tris buffer (pH 8) were added to the wells to yield final concentrations of 5 mM, 1mM and 20 µg/mL, respectively. Wherever appropriate, a TCEP (20 mM) stock prepared in Tris buffer (pH 8) was added to a final concentration of 1 mM. The plates were read using the BioTek Synergy H1 Microplate Reader. The instrument was set to run on the endpoint/kinetic luminescence setting. Luminescence was set to read every 5 minutes for the run, usually between 30 minutes to 2 hours. Luminescence Fiber was the optics type. The gain was set to 240, and the integration time was set to 0:01:00. The temperature was set to 37 °C.

**CFU Analysis**. Overnight cultures of luciferase-expressing *E. coli* were grown, and luciferase protein expression was induced as above. Subsequently, washing was performed by first pelleting the cells in the HERAEUS Multicentrifuge X1 centrifuge (Thermo Fisher Scientific) at 3300g for 2-3 minutes. The supernatant was then discarded, and the cells were resuspended in 1× PBS (volume same as original culture). This was repeated 2 times. The cells were resuspended after the last wash in 1× PBS. Resuspended cells were then incubated with the test compounds (CBT or PMBN) at the indicated concentrations for 30 minutes (PMBN) or one hour (CBT). Cells were then washed and resuspended as described above. Serial dilutions of the cells were then carried out in PBS up to a dilution factor of $10^{-8}$. 80 µL of chosen dilutions were then plated on LB agar plates with 50 µg/mL kanamycin and incubated at 37 °C for 16 hours. Subsequently, the number of colonies was manually counted.

**Nitrocefin Assay.** Overnight cultures of luciferase-expressing *E. coli* were grown, and luciferase protein expression was induced as above to mimic conditions of the cells used in the bioluminescence assay. Cells were then washed twice and resuspended in 1× PBS and then incubated with the different concentrations of PMBN (same as those concentrations used in the bioluminescence assays) with 50 µg/mL nitrocefin for 30 minutes. As a negative control, cells were treated with PBS, and as a positive control cells were treated with 10 mM EDTA for 30 minutes. At the end of the incubation period, absorbance of the cells was read at 486 nm using the BioTek Synergy H1 Microplate Reader. Untreated cells with no nitrocefin added were treated as the blank. Wherever presented, the absorbances were blank-subtracted.

**SYTOX Green Assay.** Overnight cultures of luciferase-expressing *E. coli* were grown, and luciferase protein expression was induced as above to mimic conditions of the cells used in the bioluminescence assay. Cells were then washed twice and resuspended in 1× PBS and then incubated with the test compounds (same concentrations as those used in the bioluminescence assays) for the appropriate time durations (30 minutes for PMBN and 1 hour for antibiotic conjugates). Cells were then washed twice again and resuspended in 1× PBS. The experiment was then conducted as per the manufacturer's protocol. As a negative control, cells were treated with PBS, and as a positive control cells were treated with 10 mM EDTA. Cells were analyzed using an Attune NxT flow cytometer equipped with a 488 nm laser and 525/40 nm bandpass filter. The data were analyzed using the Attune NxT Software, where populations were gated and no less than 10,000 events per sample were recorded. Wherever presented, the mean fluorescence intensity (MFI) is the ratio of fluorescence levels above the negative control treatment.

**Kinetic Model of Cytosolic Accumulation of Small Molecules in *E. coli*.** A series of kinetic equations were used to define our system, where 'X' denotes our molecule of interest, 'k' denotes the rate constants, superscripts 'out' and 'in' distinguish between exogenous and endogenous molecules, and asterisk represents luminescence:

$$\text{D-Cys-X}^{(out)} \underset{k_{-1}}{\overset{k_1}{\rightleftarrows}} \text{D-Cys-X}^{(in)} \qquad \text{(Transport of D-Cys-X at bacterial membranes)} \qquad (S1)$$

$$\text{D-Cys-X}^{(in)} \underset{k_{-2}}{\overset{k_2}{\rightleftarrows}} \text{D-Cys} + \text{X} \qquad \text{(S-S breakage resulting in release of D-Cys)} \qquad (S2)$$

$$\text{D-Cys} + \text{CBT} \overset{k_3}{\rightarrow} \text{D-luciferin} \qquad \text{(Production of D-luciferin via a click rxn)} \qquad (S3)$$

$$\text{D-luciferin} \overset{k_4}{\rightarrow} \text{D-luciferin}^{(*)} \qquad \text{(Generation of luminescence via an enzymatic rxn)} \qquad (S4)$$

Eq. S1-S4 are used to generate a series of coupled differential equations which describe the time varying evolution of concentration of each component:

$$\frac{\partial}{\partial t}[\text{D-Cys-X}^{(out)}] = -k_1[\text{D-Cys-X}^{(out)}] + k_{-1}[\text{D-Cys-X}^{(in)}] \quad (S5)$$

$$\frac{\partial}{\partial t}[\text{D-Cys-X}^{(in)}] = k_1[\text{D-Cys-X}^{(out)}] - (k_{-1}+k_2)[\text{D-Cys-X}^{(in)}] + k_{-2}[\text{D-Cys}][\text{X}] \quad (S6)$$

$$\frac{\partial}{\partial t}[\text{D-Cys}] = k_2[\text{D-Cys-X}^{(in)}] - k_3[\text{D-Cys}][\text{CBT}] - k_{-2}[\text{D-Cys}][\text{X}] \quad (S7)$$

$$\frac{\partial}{\partial t}[\text{CBT}] = -k_3[\text{D-Cys}][\text{CBT}] \quad (S8)$$

$$\frac{\partial}{\partial t}[\text{D-luciferin}] = k_3[\text{D-Cys}][\text{CBT}] - k_4[\text{D-Cys}] \quad (S9)$$

$$\frac{\partial}{\partial t}[\text{D-luciferin}^{(*)}] = k_4[\text{D-luciferin}] \quad (S10)$$

The solutions to Eq. S5-S10 yield the base components (i.e., [D-Cys-X$^{(out)}$], [D-Cys-X$^{(in)}$], [D-Cys], D-luciferin, D-luciferin$^{(*)}$) which describe the time-dependent responses observed in the luminescence experiments. Specifically, the time-resolved luminescence response can be described by Eq. S11, where $\varepsilon_0$ and $I_{baseline}$ represent the luminescence coefficient factor and baseline signal, respectively. We note that the baseline signal in our experiments pertains to those trials where only the CBT molecule was applied.

$$I_{Lumin.} = \varepsilon_0 \times [\text{D-luciferin}^{(*)}] + I_{baseline} \quad (S11)$$

Following this, the time-resolved luminescence findings serve as constraints for deriving a physically meaningful solution to Eq. S5-S10, specifically to determine rate constants. This subsequently allows for the fitting of the measured data. Fitting was done using Igor Pro (WaveMetrics, version 6.32A). During the fitting process, we established simplified, yet robust assumptions as follows:

(a) $k_{-1}$ = 0.0001 min$^{-1}$, as the overall accumulation becomes defined by $k_1$.

(b) $k_{-2}$ = 0.0001 M$^{-1}$.min$^{-1}$, as the next step is a very fast process.

(c) $k_3$ = 156 M$^{-1}$.min$^{-1}$, as reported in literature[1]

(d) $k_4$ = 96 min$^{-1}$, as reported in literature[2]

**Synthesis of Antibiotic Conjugates.**

**General Methods.** Reverse phase preparative high-performance liquid chromatography RP-HPLC purification was performed on instruments equipped with Waters 1525 pumps and 2489 UV/Visible Detector on a Phenomenex Luna 10 µm C8(2) 100 Å (250 x 21.2 mm) or C18 columns using a 5 to 100% linear gradient of methanol (MeOH) in $H_2O$ or acetonitrile (MeCN) in water each containing 0.1% TFA at 10 mL/min. The HPLC fractions of the desired compounds were first concentrated under reduced pressure using a rotary evaporator. The concentrated aqueous solutions were then lyophilized with Labconco Freezone 4.5L lyophilizer (- 84 °C). The purity of the samples was ascertained by analytical RP-HPLC using a Phenomenex Luna 5 µm C8(2) 100 Å (250 x 4.6 mm) on the same instrument; using gradient elution in $H_2O$/MeCN or $H_2O$ /MeOH with 0.01% TFA in each solvent at 1 mL/min prior to use in biological assays. Electrospray ionization (ESI)-based mass analysis was performed on Advion Expression® CMS mass spectrometer using standard parameters for intermediates. A low fragmentation, low energy setup was used to analyze fragmentation-sensitive compounds. High resolution mass spectrometry (HRMS) analysis of the compounds was done using a C18(2) column (Luna 5 µm 100Å 250 x 4.6 mm) connected to an Agilent LC-QTOF (Agilent 1260 Infinity II Prime LC with Agilent 6545B QTOF). The mass of the peptides was analyzed by MS using MassHunter software. UV spectroscopic analysis was performed on Genesys 50 (Thermo Scientific) UV-Visible Spectrophotometer.$^{1}$H and $^{13}$C-NMR spectra for final compounds and intermediates were acquired on a Varian 600MHz spectrophotometer. All NMR spectra were processed and analyzed using MestreNova software. Residual solvent signals from deuterated solvents were used as an internal standard with reference to tetramethylsilane (TMS) for defining chemical shifts. Chemical shifts are reported in ppm (δ) and coupling constants (J) are reported in Hertz [Hz].

**Ciprofloxacin-methyl-ester (cys-Cipro) Conjugate.**

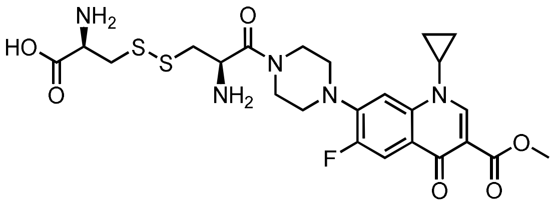


**Molecular Weight: 567.65140**

**Synthesis of Ciprofloxacin-methyl-ester.** Synthesis was performed as per a previously reported procedure in literature.[3] Column chromatography on silica gel (Supelco, 60 Å, 230-400 mesh, 40-63 m particle size) was performed on manually packed columns with rationally chosen solvents. The compound was then characterized by mass and $^{1}$H-NMR spectroscopy.

MF: $C_{19}H_{22}FN_3O_3$, calcd. m/z 346.15 obs. 346.1 for $[M+H]^+$. $^{1}$H-NMR: ($CDCl_3$) d 1.14 (m, 2H, cyclopropyl-$CH_2$), 1.32 (m, 2H, cyclopropyl-$CH_2$), 3.12 (m, 4H, piperidyl 2 x $CH_2$), 3.27 (m, 4H, piperidyl 2 x $CH_2$), 3.43 (m, 1H, cyclopropyl CH-N), 3.91 (s, 3H, $OCH_3$), 7.26 (d, *J* = 6 Hz, 1H, ArH), 8.02 (d, *J* = 18 Hz, 1H, ArH), 8.54 (s, 1H, ArH).

**Synthesis of D-Cystine-Ciprofloxacin-methyl-ester.** Commercially available N-$^{t}$Boc-D-Cys was oxidized to N-di-$^{t}$Boc-D-Cystine by dissolving 200 mg (0.905 mmol) of compound in a solution containing 22.5 mL MeCN and 7.5 mL 0.5 M ammonium bicarbonate. The solution was bubbled with air overnight while stirring at room temperature (~18hrs). An Ellman's test of the reaction mixture revealed no remaining free thiols. At this point, the reaction mixture was concentrated under reduced pressure using a rotary evaporator and lyophilized to dryness, yielding 185 mg of disulfide (97%), which was used after removal of ammonium bicarbonate. Conjugation of N-di-$^{t}$Boc-D-Cystine to ciprofloxacin-methyl-ester was performed as follows: to a solution of N-di-$^{t}$Boc-

D-Cystine (135 mg, 0.307mmol) in 10-15 mL dimethylformamide (DMF), cipro-methyl-ester (50mg, 0.145mmol), 4-dimethylaminopyridine (DMAP, 4mg, 0.03mmol,), and 1-ethyl-3-(3-dimethylaminopropyl) carbodiimide (EDC, 25 mg, 0.130mmol) were added. The mixture was stirred at room temperature. Thin layer chromatography (TLC) and electrospray ionization (ESI) mass spectroscopy were used to monitor the reaction progress. After 24 hrs of stirring, the mixture was subjected to liquid-liquid phase extraction using water and dichloromethane (DCM) to remove DMF and other water-soluble byproducts. The organic layer was separated, washed with 0.1 M hydrochloric acid (HCl) and dried over $Na_2SO_4$ to be then concentrated under vacuo to obtain crude residue. Column chromatography of the crude residue was then carried out on manually packed columns over silica-gel (Supelco, 60 Å, 230-400 mesh, 40-63 m particle size) using chloroform first and then an increasing gradient of chloroform: methanol. Appropriate fractions were collected and combined which yielded a mixture of N-di-$^t$Boc-D-Cystine conjugated to either one or two molecules of ciprofloxacin-methyl-ester. These fractions were concentrated and then subjected to deprotection of the $^t$Boc group by treatment with 0.5 mL TFA + 0.5 mL DCM with stirring at room temperature for 2 hours. The reaction mixture was concentrated using the rotary evaporator and triturated with diethyl ether to remove TFA, to obtain a crude solid. This crude solid was then purified using RP-HPLC. Fractions containing m/z of desired product were collected and concentrated under reduced pressure using a rotary evaporator, to yield a transparent residue, which was lyophilized to a white solid (21mg). In later fractions, the expected bis ciprofloxacin side-product was observed in minor amounts. The desired compound was characterized with $^1$H-NMR and mass spectrometry prior to being used in permeability assays. A stock solution of the compound was prepared by dissolving in Millipore-DI/$H_2O$. The stock solution was then analyzed for purity by analytical RP-HPLC. Peaks were broadened in NMR due to H/D exchange.

$^1$H-NMR: ($D_2O$) $\delta$ 1.14 (m, 2H, cyclopropyl-$CH_2$), 1.32 (m, 2H, cyclopropyl-$CH_2$), 1.39-1.49 (m, 6H, 3 x $CH_2$), 1.49 (s, 9H, 3 x$CH_3$), 1.58 (dt, *J*=6Hz, 128Hz, 2H, $CH_2$), 1.74 (dt, *J*=6, 12Hz, 2H, $CH_2$), 3.22 (t, , *J*=6Hz, 4H, 2 x $CH_2$), 3.43 (m, 1H, CH), 3.45-3.51 (m, 4H, piperidyl 2 x CH2), 3.62 (m, 2H), 3.63-3.67 (m, 4H, 2 x $CH_2$), 7.31 (d, *J* = 6 Hz, 1H, ArH), 8.03 (d, *J* = 12 Hz, 1H, ArH), 8.81 (s, 1H, ArH), 10.1 (brs, 1H, NH). $^{13}$C-NMR: ($CDCl_3$) d 8.2, 25.4, 26.8, 28.4, 29.5, 32.6, 34.7, 39.2, 45.1, 50.0, 70.0, 70.2, 70.6, 71.3, 105.0, 111.4, 112.7, 112.8, 122.2, 138.4, 144.8, 146.8, 152.6, 154.2, 154.6, 165.1, 175.4. HRMS-QTOF: for MF; $C_{24}H_{31}FN5_4O_6S_2$ calcd. m/z 568.1694 obs. 568.1699 for $[M+H]^+$

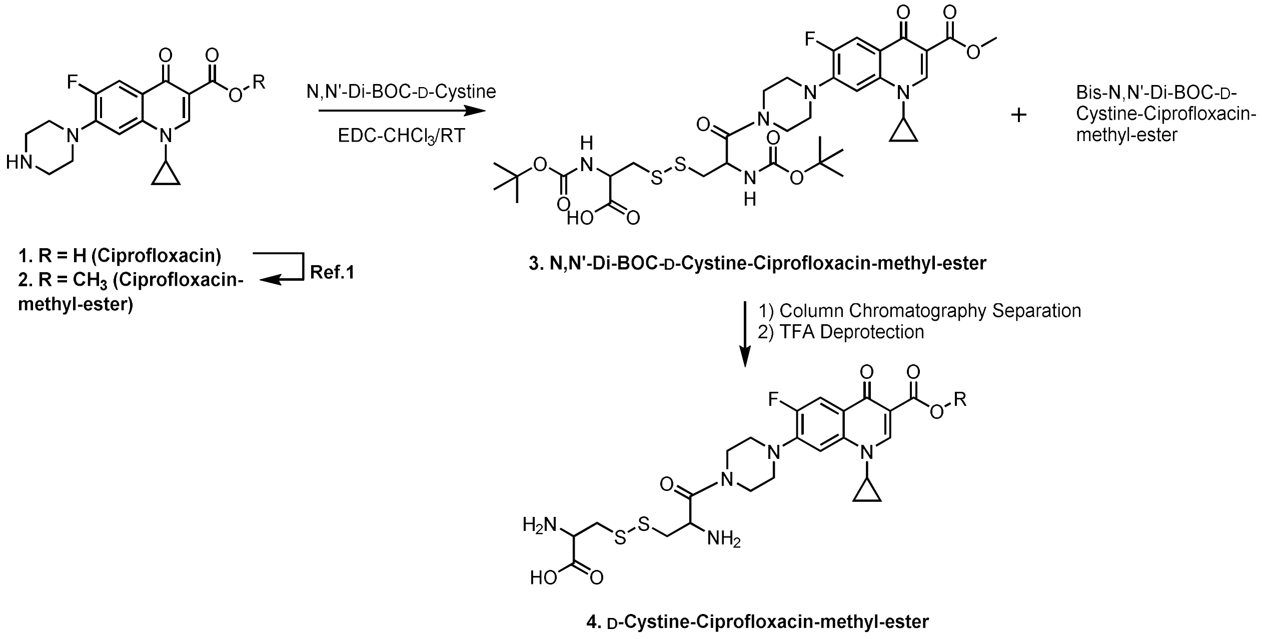


Synthesis Scheme for **cys-Cipro** or **D-Cystine-Ciprofloxacin-methyl-ester.**

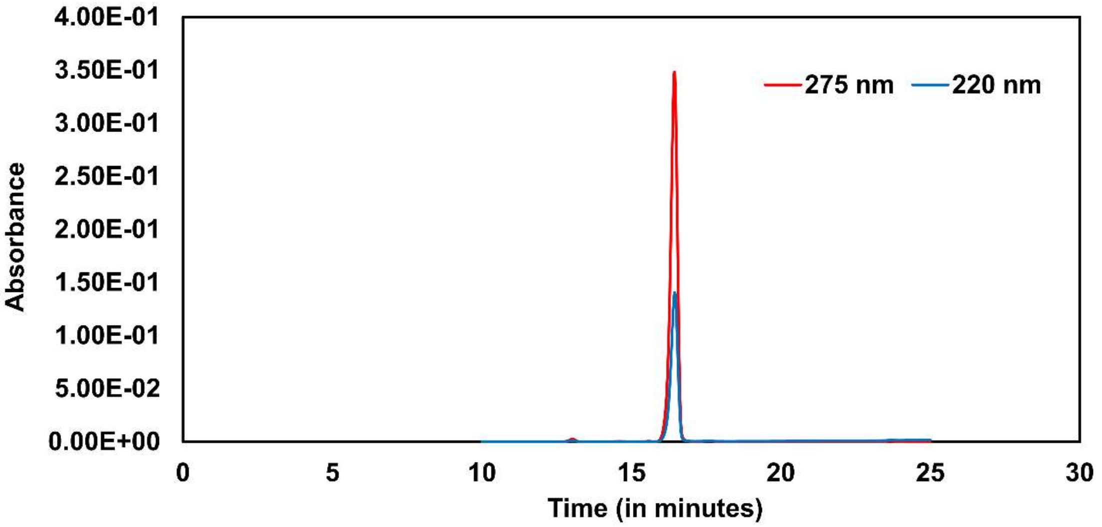


Analytical HPLC Chromatogram of **cys-Cipro** or **D-Cystine-Ciprofloxacin-methyl-ester.**

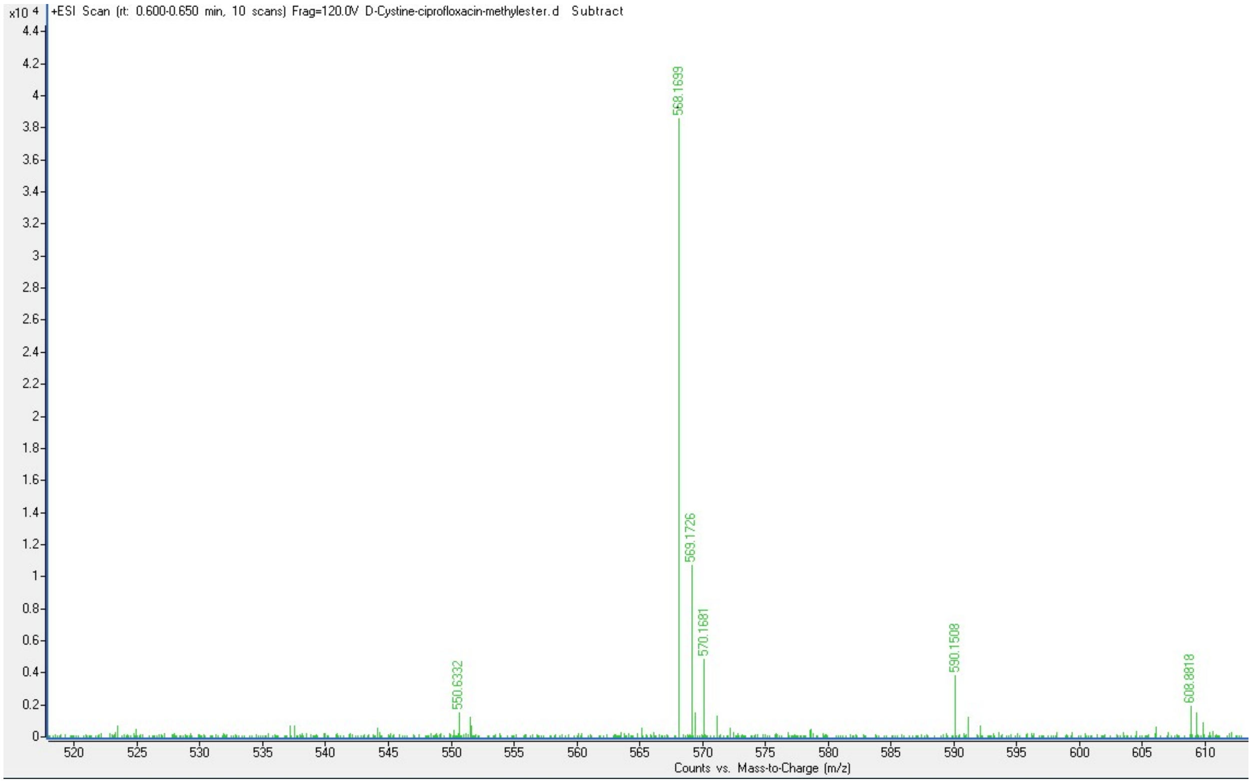


QTOF High Resolution Mass Spectrum for **cys-Cipro** or **D-Cystine-Ciprofloxacin-methyl-ester** (m/z 568 for $[M+H]^+$).

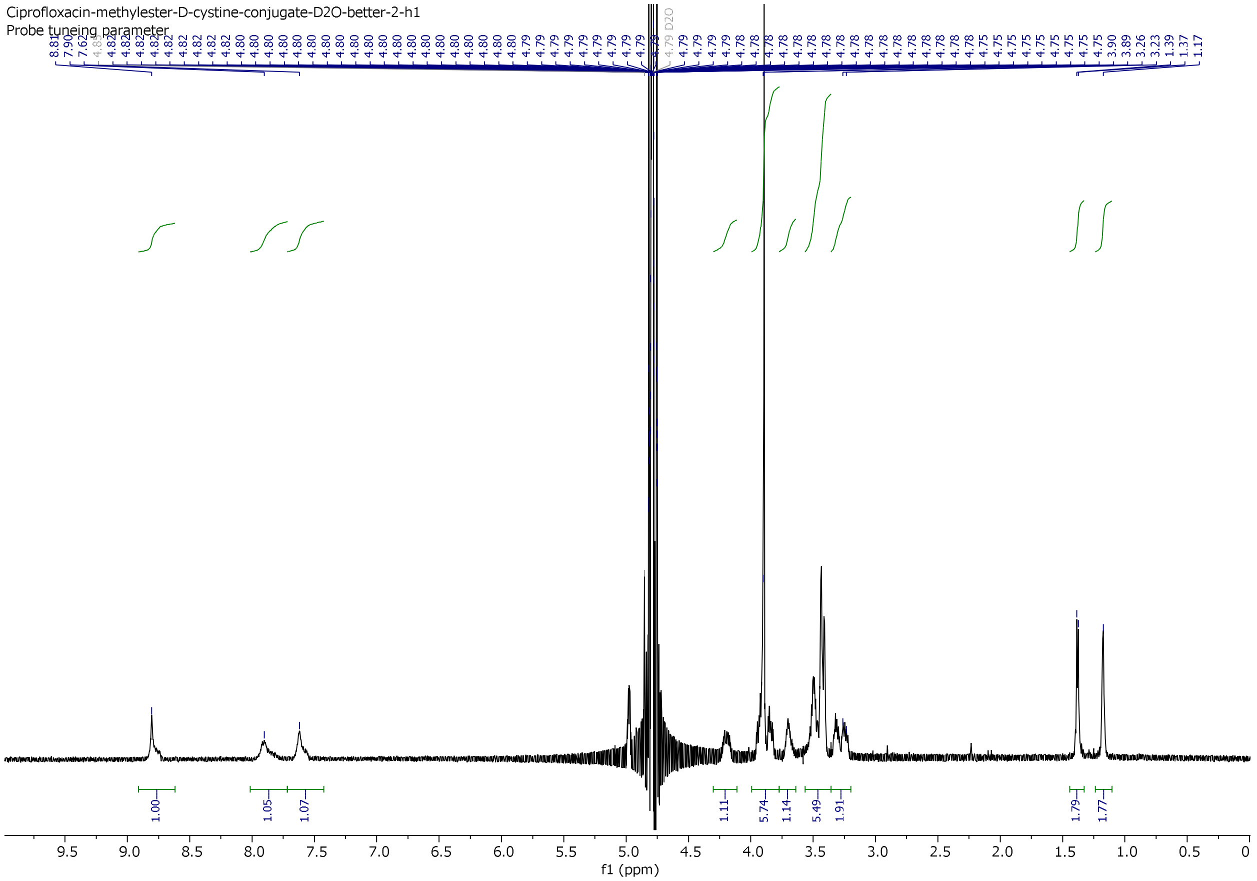


[1]H NMR spectrum of **cys-Cipro** or **D-Cystine-Ciprofloxacin-methyl-ester.**

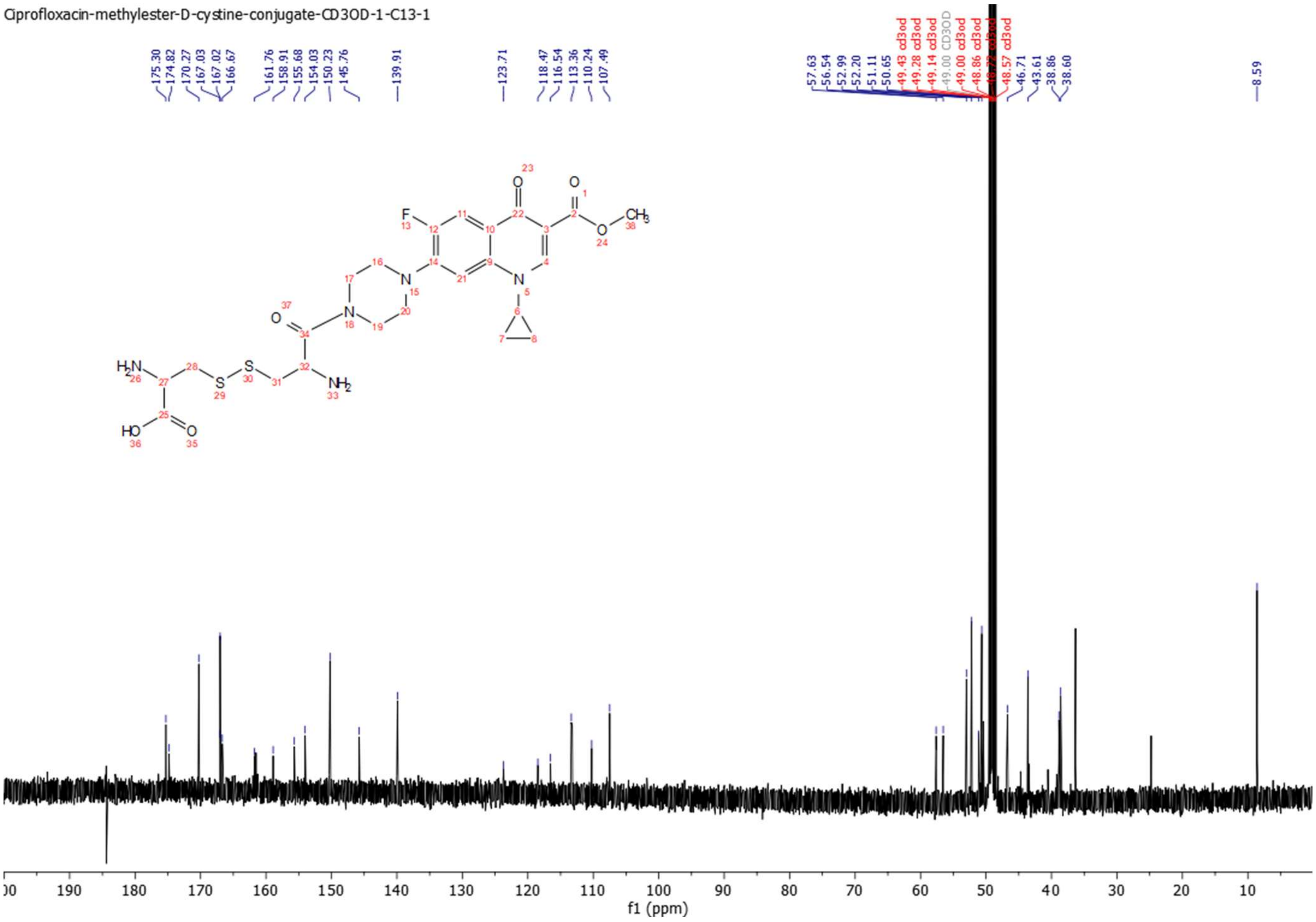


$^{13}C$ NMR Spectrum of **cys-Cipro** or **D-Cystine-Ciprofloxacin-methyl-ester.**

**Linezolid (cys-Line) Conjugate.**

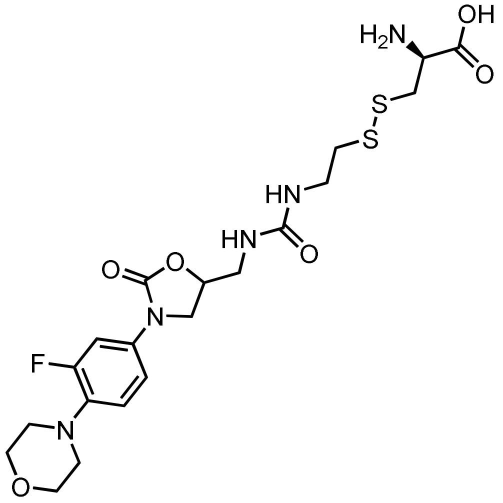

**Molecular Weight: 517.59140**

**Synthesis of Linezolid-Cystamine-D-Cys-Disulfide.** Synthesis of the compound was performed as shown in the synthetic scheme below, linezolid amine (30.0 mg, 0.1 mmol) was dissolved in 3 mL of anhydrous $CH_2Cl_2$ and carbonyldiImidazole (18 mg, 0.11 mmol) as well as DIEA (10 µL) was added to it. The homogenous mixture was stirred at room temperature overnight. Next day, the mixture was concentrated under reduced pressure and a solution of cystamine-pyridine disulfide-hydrochloride (23.5 mg, 0.1 mmol) in 3 mL anhydrous MeCN and DIEA (20 µL) was added to it. The mixture was stirred at room temperature for 4 hours, during which ESI mass analysis indicated that the starting material was consumed to form linezolid-pyridyl-cystamine disulfide conjugate (m/z was observed). The mixture was then concentrated under reduced pressure on the rotary evaporator, and the leftover residue was redissolved in methanol. To it, an aqueous solution of D-Cys (25.0mg, 0.2mmol, in 2.0 mL DI water) was added. Upon stirring the solution for 2 hours at room temperature, ESI mass analysis indicated the formation of the desired linezolid-cystamine-D-Cys-disulfide (m/z was observed). The reaction mixture was concentrated under reduced pressure followed by dissolution in MeCN: $H_2O$ (1:3, 10 mL). This solution was then purified using RP-HPLC. Fractions containing m/z of desired product were collected and concentrated under reduced pressure using a rotary evaporator to yield a colorless solid (27.7 mg, 53%). The desired compound was characterized with $^{1}$H-NMR, $^{13}$C-NMR and mass spectrometry prior to being used in permeability assays. A stock solution of the compound was prepared by dissolving in Millipore-DI/$H_2O$. The stock solution was then analyzed for purity by analytical RP-HPLC

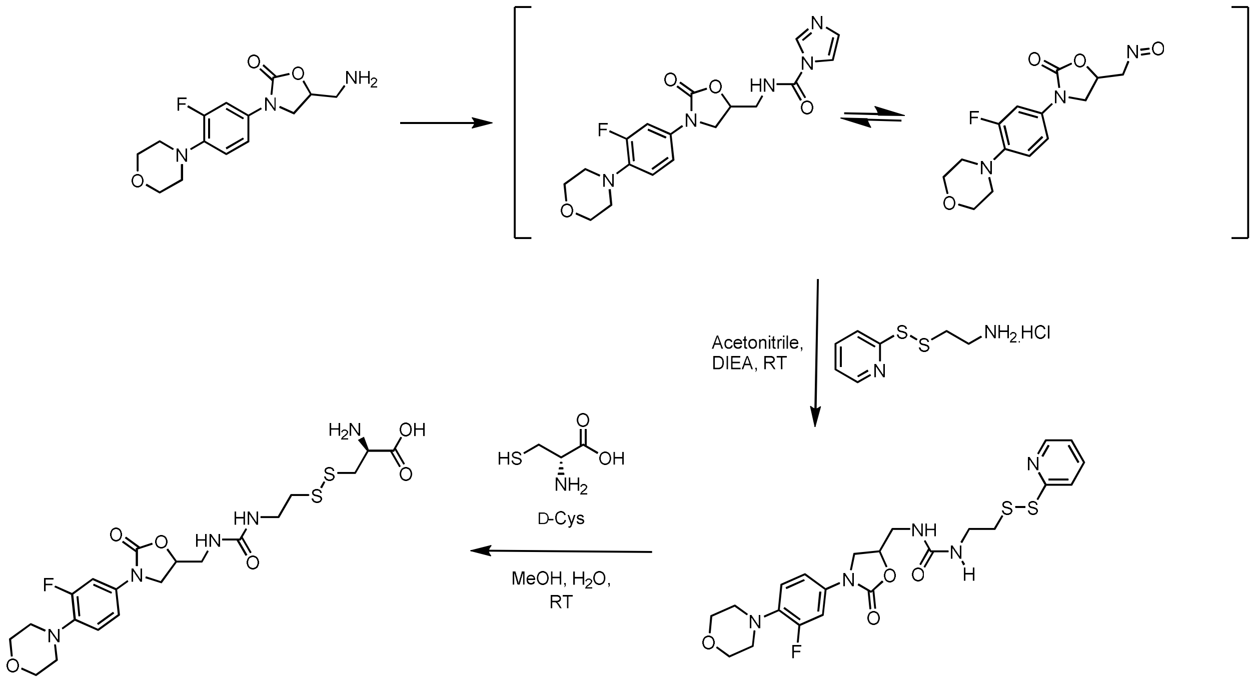


Synthesis Scheme for **cys-Line** or **Linezolid-Cystamine-D-Cys-Disulfide.**

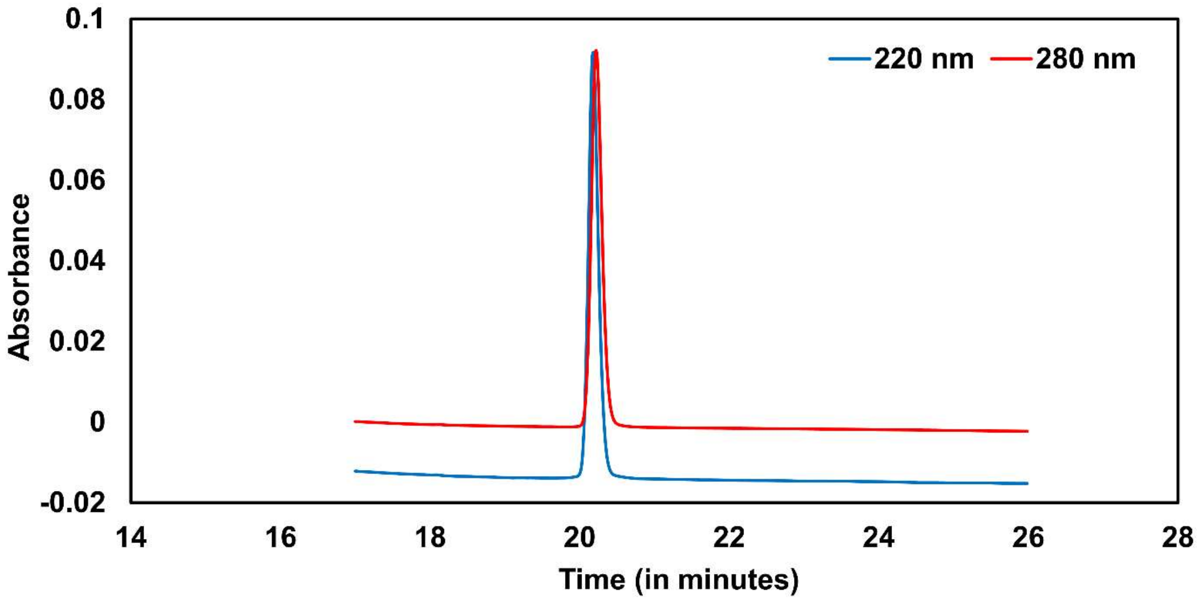


Analytical HPLC Chromatogram of **cys-Line** or **Linezolid-Cystamine-D-Cys-Disulfide.**

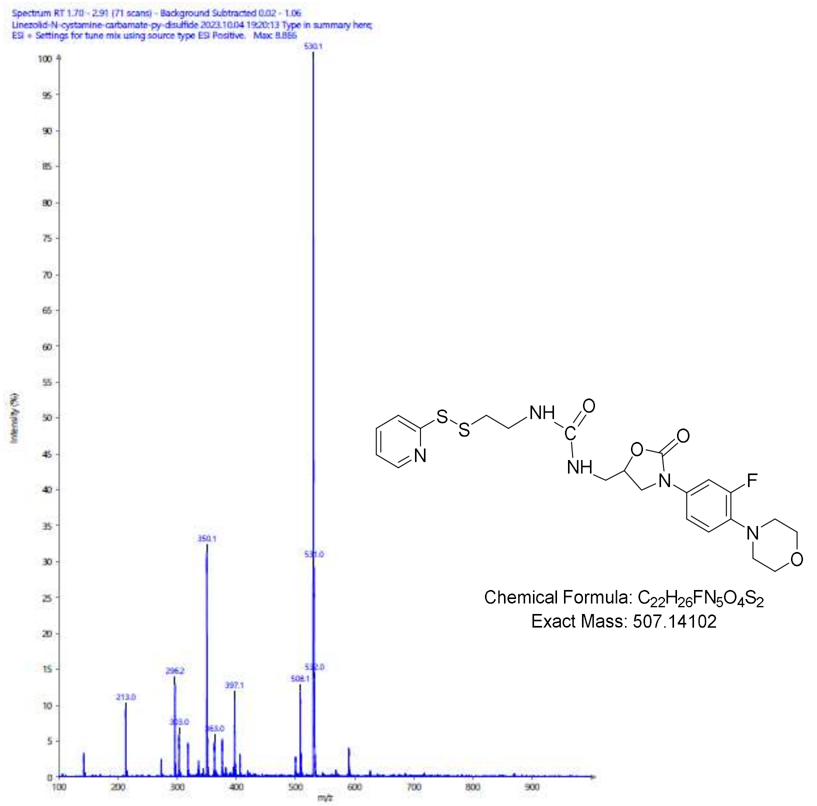


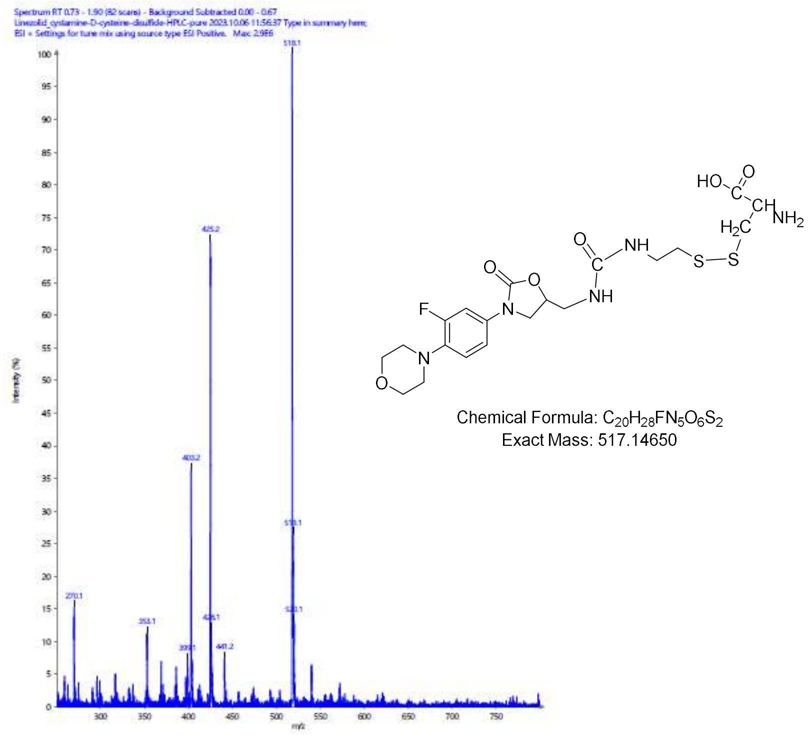


ESI-Mass Spectra for intermediate (m/z 508 for $[M+H]^+$) and **cys-Line** or **Linezolid Cystamine-D-Cys-Disulfide** (m/z 518 for $[M+H]^+$).

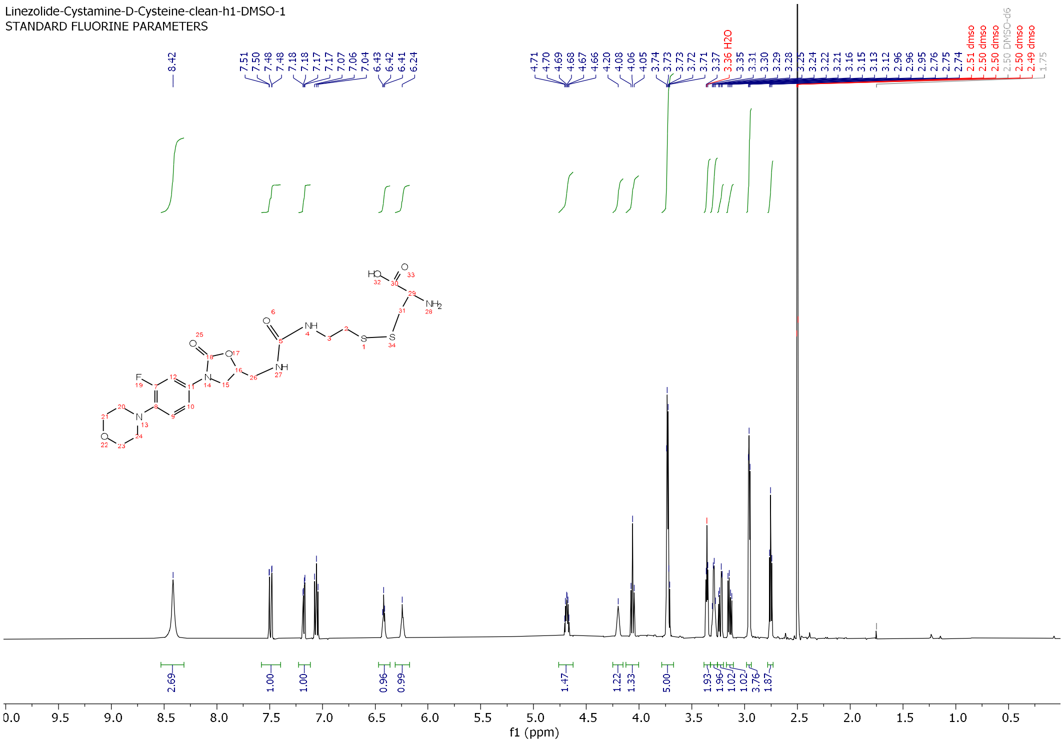


$^1$H NMR spectrum of **cys-Line** or **Linezolid-Cystamine-D-Cys-Disulfide.**

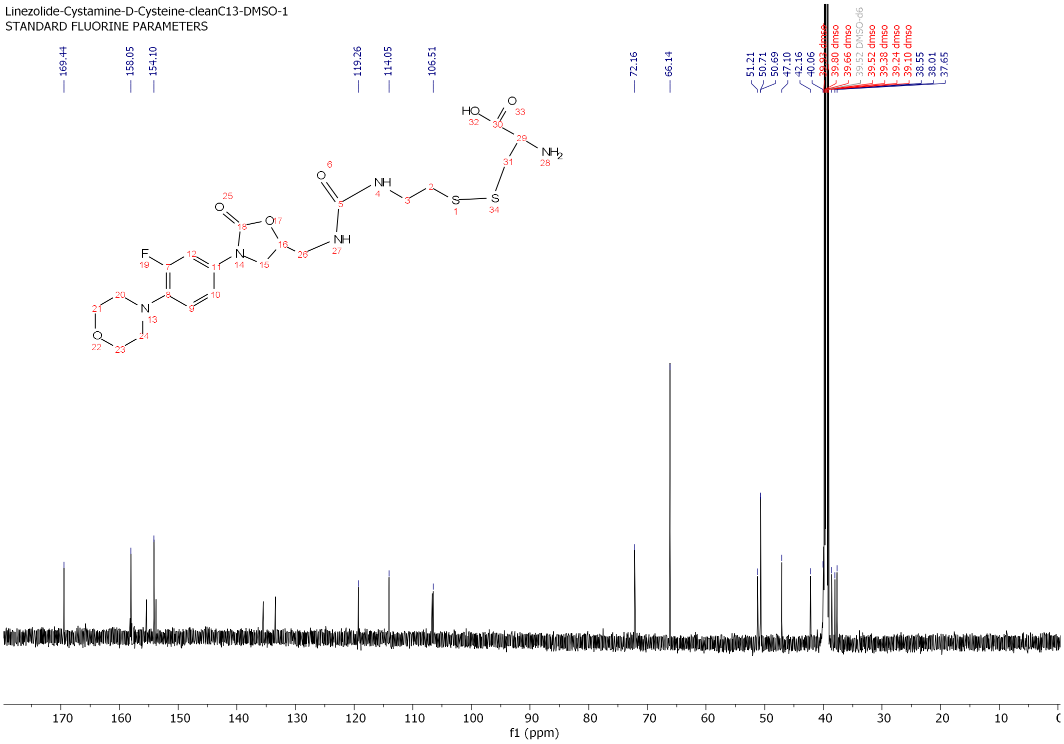


$^{13}$C NMR Spectrum of **cys-Line** or **Linezolid-Cystamine-D-Cys-Disulfide.**

**Puromycin (cys-Puro) Conjugate.**

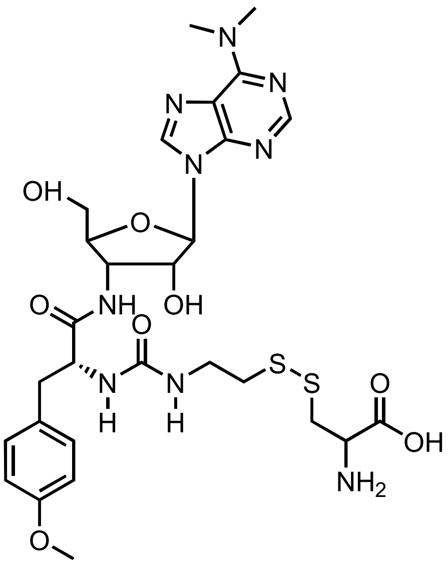


Molecular Weight: 693.79500

**Synthesis of Puromycin-Cystamine-D-Cys-Disulfide.** Synthesis was performed as shown in synthetic scheme below. Puromycin dihydrochloride (procured from Alfa Aesar, Cat # J61278) (27.4mg, 0.05mmol) was dissolved in 2 mL of anhydrous DMF and to it was added preformed 2-((2-isocyanatoethyl)disulfaneyl)pyridine (20.0mg, from carbonyldiimidazole and 2-(pyridine-2-yldisulfaneyl)ethan-1-amine), followed by DIEA (10uL). The homogenous mixture was stirred at room temperature overnight. Next day, upon observation by ESI mass analysis that the intermediate pyridyl disulfide was formed, the mixture was triturated with ether (3 x 5 mL) to remove DMF as much as possible. The leftover reside was redissolved in methanol and to it was added an aqueous solution of D-Cys (75.0mg, 0.4mmol, in 2.0 mL DI water). Upon stirring the mixture for 2 hr at room temperature, ESI mass analysis indicated the desired puromycin-cystamine-D-Cys-disulfide was formed (m/z 694 for $[M+H]^+$). The reaction mixture was concentrated under reduced pressure to remove methanol. The leftover solution was dissolved in MeCN: $H_2O$ (1:4, 10 mL) and the solution was subjected to preparative RP-HPLC purification on a C18-silca column. The homogenous fractions containing desired compound as analyzed by ESI-mass were combined and concentrated to remove MeCN, the remaining solution was frozen to -80 °C for 2 hours, and then the frozen sample lyophilized to yield colorless solid (23.8 mg). The compound was isolated as a trifluoroacetate salt and was characterized by mass spectrometry, $^1$H-NMR spectroscopy. The stock solution was then analyzed for purity by analytical RP-HPLC. The spectroscopic data was in agreement with the assigned structure.

$^1$H-NMR: (DMSO-$d_6$) d 2.69-2.74 (m, 3H, -$CH_2$), 2.86 (dd, 1H, *J* = 6 and 12 Hz, CH2), 3.13 (dd, 1H, *J* = 6 and 12 Hz, -CH2), 3.20-3.25 (m, 2H, -CH2), 3.46 (dd, 1H, *J* = 2 and 6 Hz, -CH), 3.69 (dd, 1H, *J* = 6 and 18 Hz, -$CH_2$), 3.92 (m, 1H), 4.19 (brs, 1H), 4.42-4.85 (m, 2H), 4.98 (d, *J* = 2 Hz, 1H), 6.21 (d, *J* = 12 Hz, 1H), 6.33 (brs, 1H), 6.82 (d, *J* = 12 Hz, 2H, ArH), 7.11 (d, *J* = 12 Hz, 2H, ArH), 8.11 (d, 1H, *J* = 6 Hz, ArH), 8.25 (s, ArH), 8.41 (brs, 1H) 8.45 (s, 1H). HRMS-QTOF: for MF; $C_{24}H_{31}FN5_4O_6S_2$ calcd. m/z 568.1694 obs. 568.1699 for $[M+H]^+$

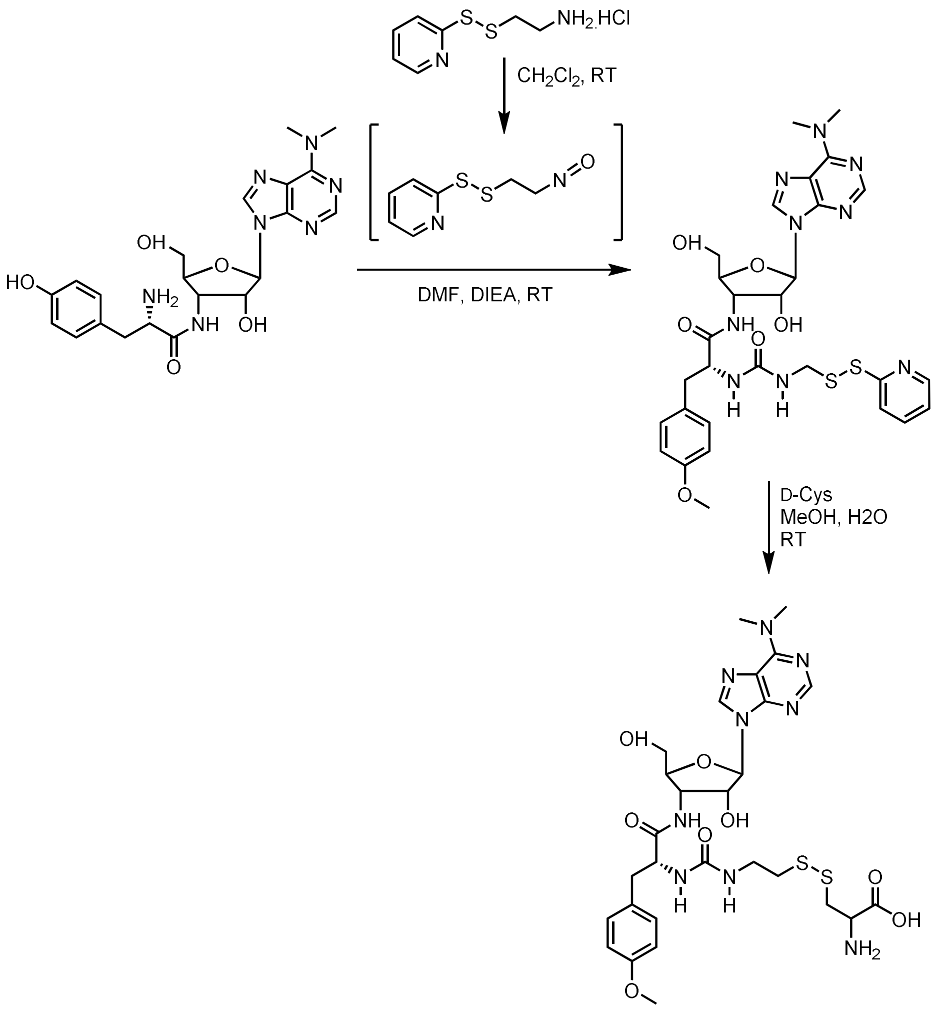


Synthesis Scheme for **cys-Puro** or **Puromycin-Cystamine-D-Cys-Disulfide.**

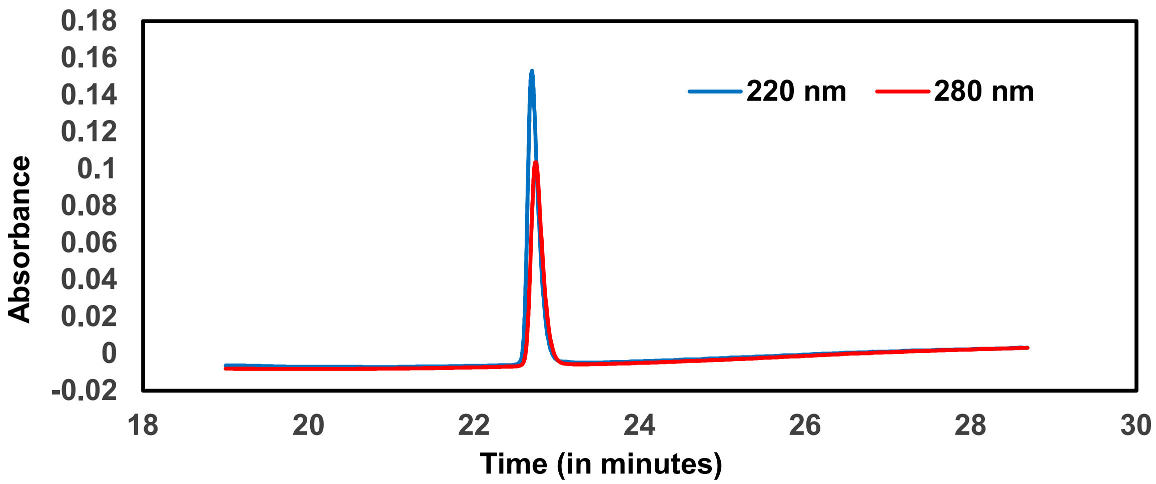


Analytical HPLC Chromatogram of **cys-Puro** or **Puromycin-Cystamine-D-Cys-Disulfide.**

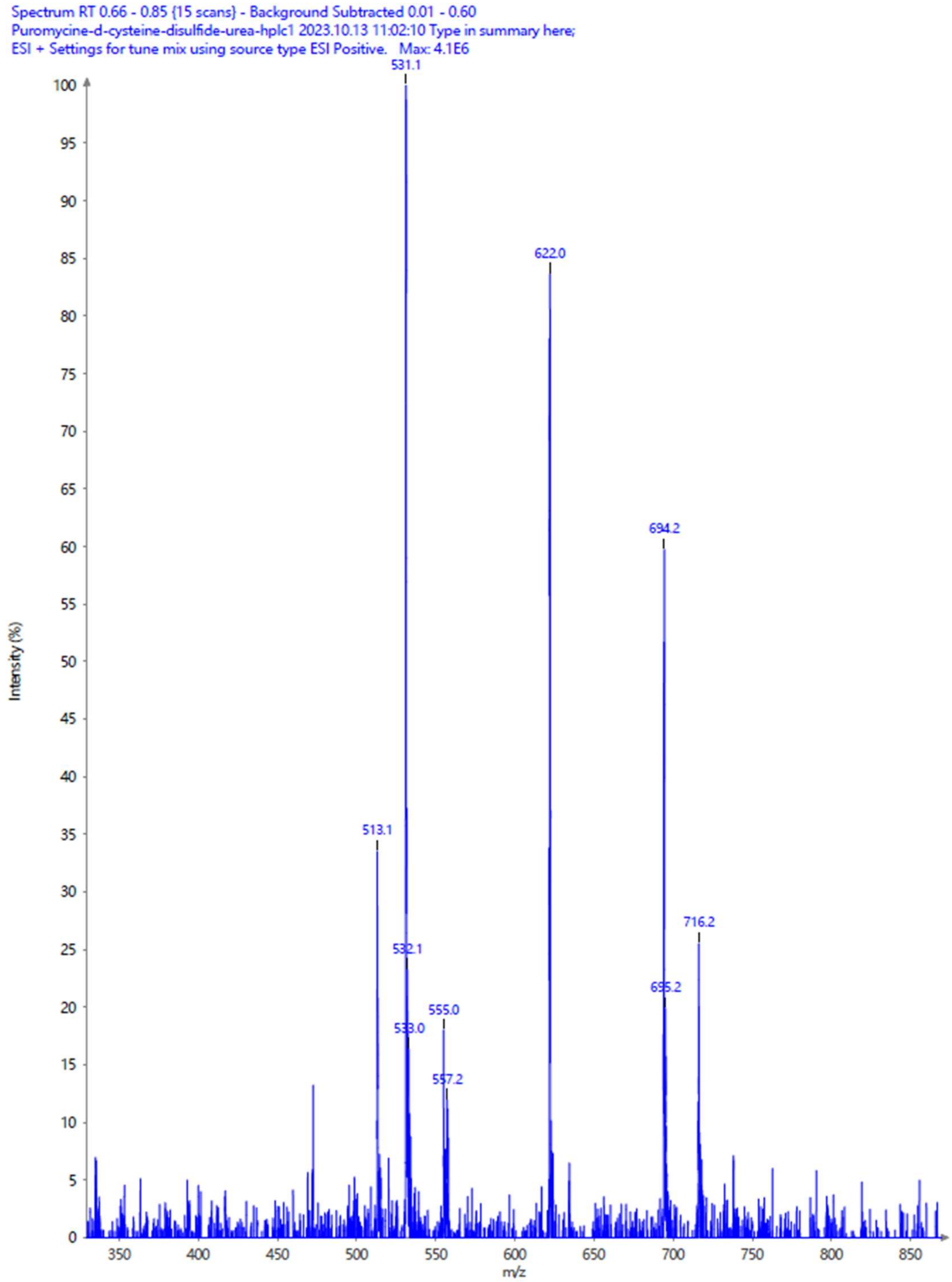


ESI-Mass Spectrum for **cys-Puro** or **Puromycin Cystamine-D-Cys-Disulfide** (m/z 694 for $[M+H]^+$).

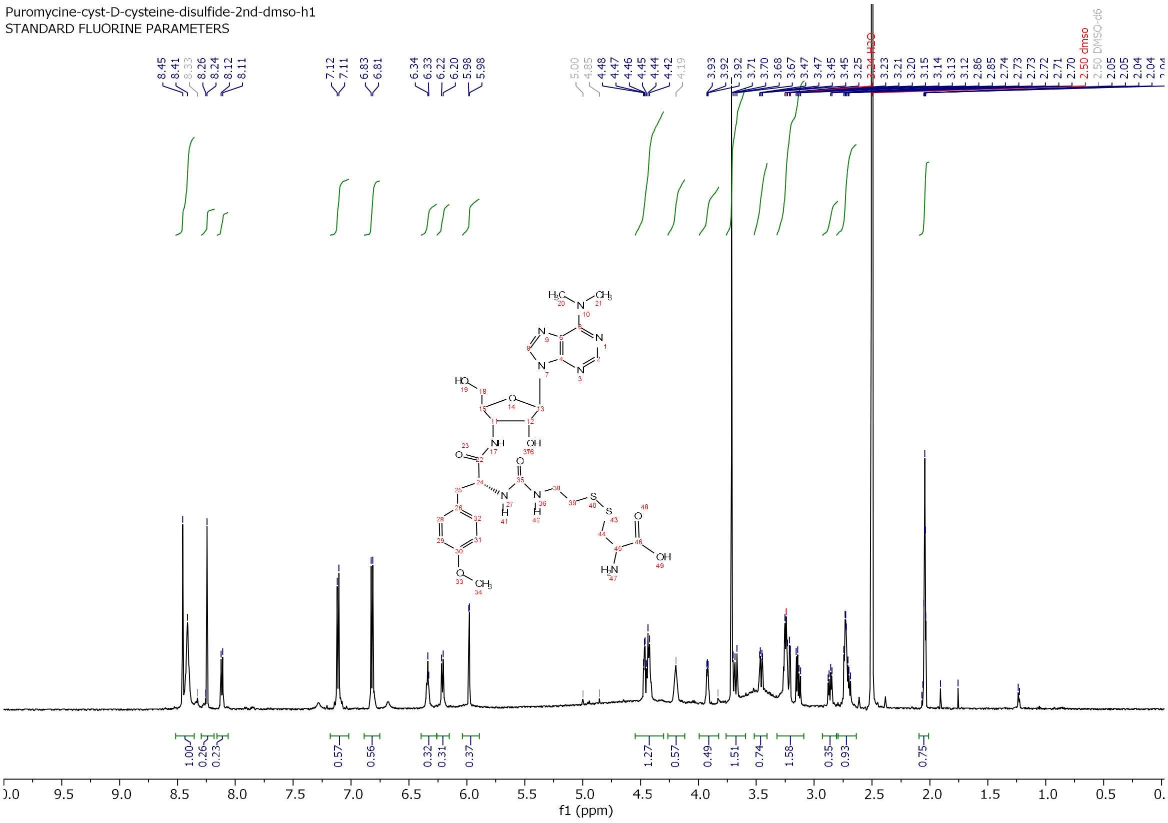


$^1$H NMR spectrum of **cys-Puro** or **Puromycin-Cystamine-D-Cys-Disulfide.**

**Rifamycin B (cys-Rifa) Conjugate.**

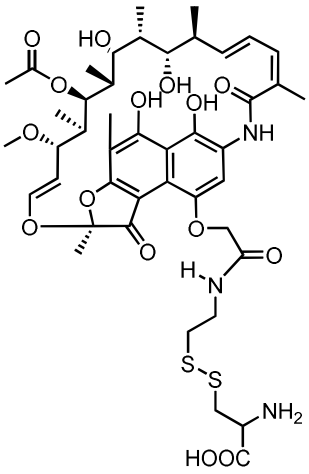

Molecular Weight: 934.08200

**Synthesis of Rifamycin-Cystamine-D-Cys-Disulfide.** Synthesis was performed as shown in synthetic scheme below, accordingly rifamycin B (25.1 mg, 0.033mmol) (procured from Toronto Research Chemicals, Cat # R508170) was dissolved in anhydrous THF (3.0cmL). To this solution,

2-(pyridine-2-yldisulfaneyl)ethan-1-amine) (11 mg, 0.05mmol), dicyclohexyl carbodiimide (DIC, 10.4mg, 0.05mmol), triethyl amine (10 uL) were added. The reaction mixture was stirred at room temperature overnight. Next day, ESI mass spectrometry analysis revealed that the starting material was consumed to form the intermediate pyridyl-disulfide (m/z 925 for $[M+H]^+$). The volatiles were removed under reduced pressure using rotary evaporator and the remaining residue was triturated with ether (2 x 5 mL). The ether layer was discarded and the left-over solid was redissolved in methanol (2 mL). To this solution was added an aqueous solution of D-Cys (25.0mg, 0.2mmol, in 1.0 mL DI water). Upon stirring the solution for 2 hr. at room temperature, ESI mass analysis indicated the desired rifamycin B-cystamine-D-Cys-disulfide (m/z 934 for $[M+H]^+$). The reaction mixture was concentrated under reduced pressure to remove methanol and the left-over was dissolved in MeCN: H2O (1:4, 10 mL). The solution was then subjected to preparative RP-HPLC purification on a C18-silca column. The homogenous fractions containing desired compound as analyzed by ESI-mass were combined, concentrated to remove as much MeCN as possible, and the leftover solution was frozen to -80 °C and then lyophilized to yield a colorless solid (20.1mg). The compound was isolated as a trifluoroacetate salt and was characterized by mass spectrometry. The stock solution was then analyzed for purity by analytical RP-HPLC. The spectroscopic data was in agreement with the assigned structure.

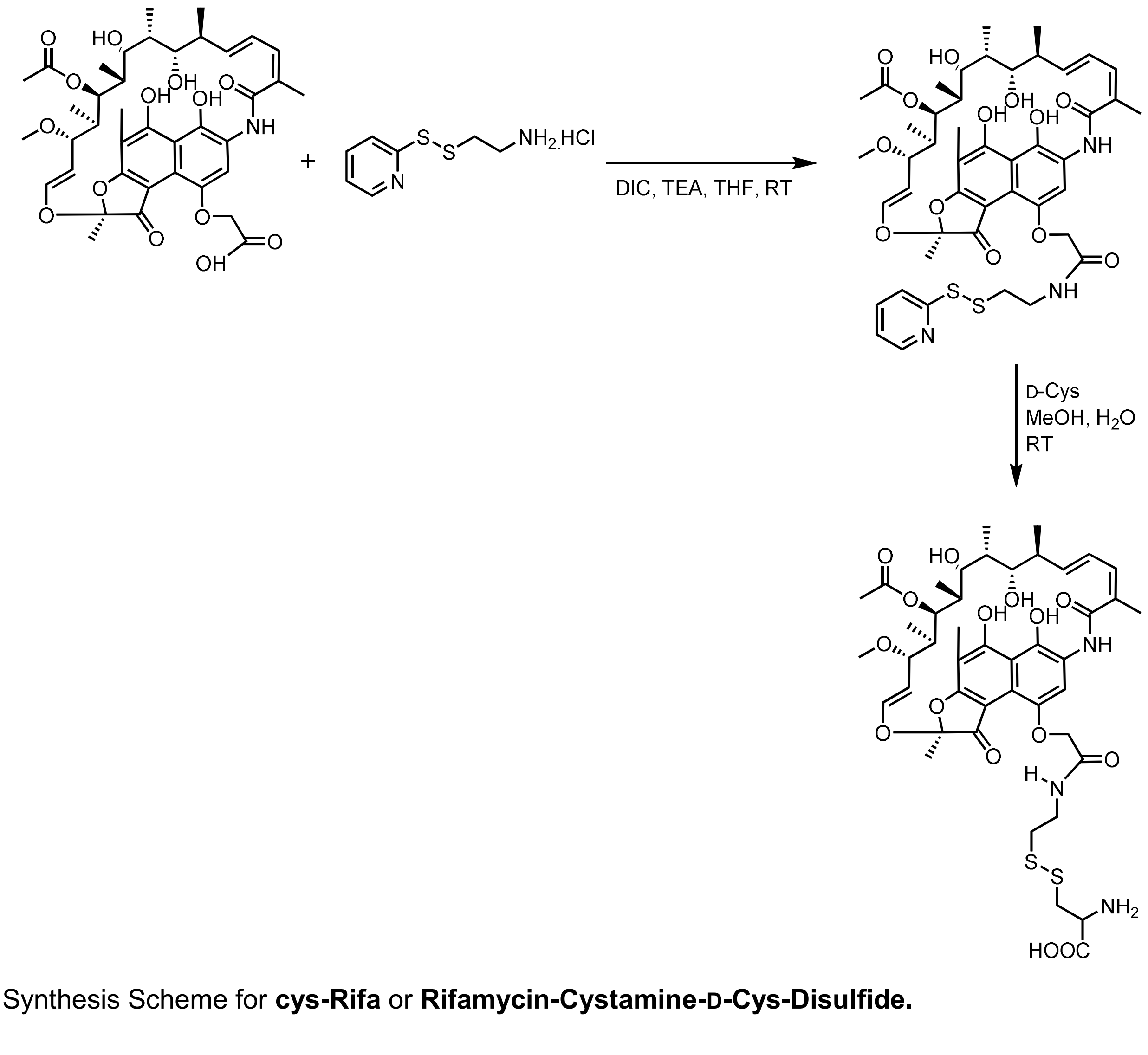


Synthesis Scheme for **cys-Rifa** or **Rifamycin-Cystamine-D-Cys-Disulfide.**

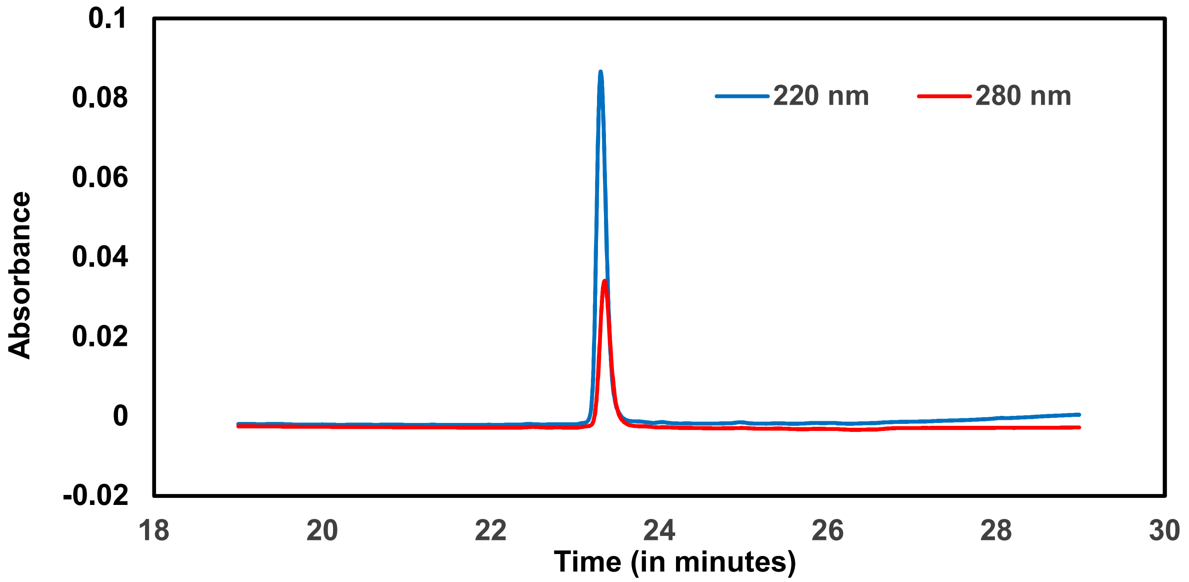


Analytical HPLC Chromatogram of **cys-Rifa** or **Rifamycin-Cystamine-D-Cys-Disulfide.**

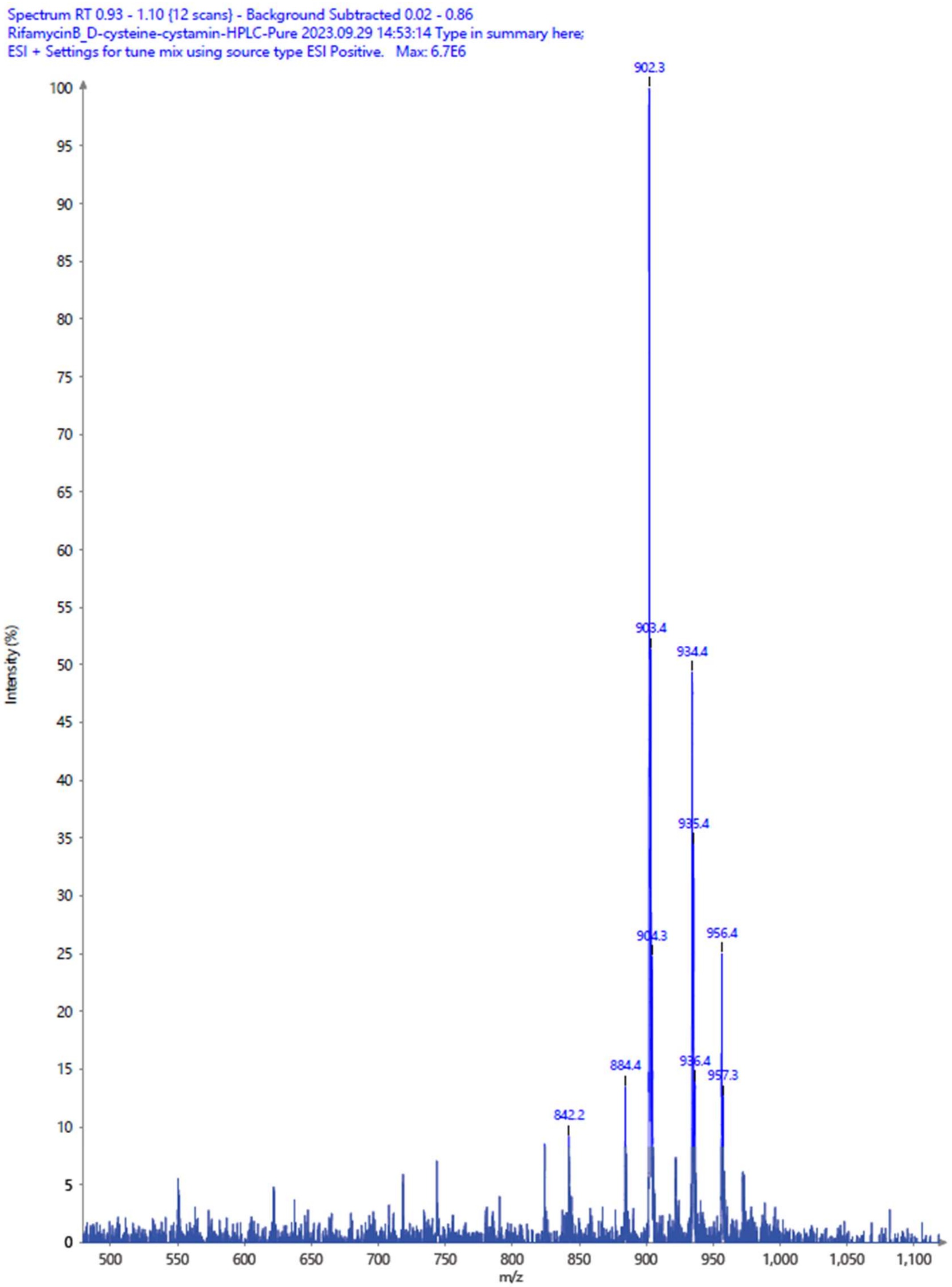


ESI-Mass Spectrum for **cys-Rifa** or **Rifamycin Cystamine-D-Cys-Disulfide** (m/z 934 for $[M+H]^+$).
